# Identification of molecular mechanisms contributing to acquired thermotolerance by transcriptome profiling

**DOI:** 10.64898/2025.12.01.691713

**Authors:** Ge Gao, Yong Woo, Christoph Gehring, Mark Tester, Magdalena M. Julkowska

## Abstract

Heat stress poses a serious threat to plant survival and productivity, and has a direct influence on crop yield stability. Plants response to high temperature is tightly controlled by complex genetic networks. Plants can be acclimated through gradual pre-exposure to increasing temperatures and that in turn causes higher survival in subsequent and otherwise lethal heat stress conditions. To investigate the physiological and molecular processes underlying heat acclimation and recovery, we examined changes in *Arabidopsis thaliana* transcriptome throughout the acclimation and the subsequent heat shock treatment. Groups of differentially expressed genes and enriched biological pathways that constitute the heat transcriptional memory were identified. The function of flavonoids in plant heat stress were further explored experimentally. In addition, we observed altered stomata density and aperture responses in heat acclimated plants, and this might be partially controlled by *AGAMOUS-LIKE16* (*AGL16*) transcription factor and its negative regulator *microRNA824* (*miR824*).

## Introduction

Plants grow in dynamic environments, where they are exposed to a plethora of stressful factors affecting their growth, development and reproductive success. An important environmental factor that often threatens plants is heat. Due to climate change, plants are facing increasing and more frequent extreme temperatures. However, plants have evolved mechanisms response to fluctuating temperatures consisting of cellular, physiological, and developmental changes to optimize their growth and reproductive success.

Plants have an inherent ability to survive certain levels of heat stress, this is called basal thermotolerance, a characteristic that varies between species and genotypes. It is also well known that plants can acquire thermotolerance when exposed to milder, sub-lethal levels of heat stress (Vierling, 1991). This pre-exposure to stress, called priming or acclimation, induces the configuration of a new cellular state that is different from the pre-stress naive state and enhances plant survival in a subsequent event of heat stress. The intervening time, during which plants experience a non-stress situation, is called the ‘memory phase” (Sedaghatmehr et al., 2016). This ability of plants to retain information from previous exposure to stress has also been found for cold, drought (Ding et al., 2012), and salinity stress (Sani et al., 2013).

Previous investigations focused on transcriptional changes in Arabidopsis either during the acclimation process (Larkindale and Vierling, 2008; Stief et al., 2014), or the memory phase (Sedaghatmehr et al., 2016). However, the processes of heat acclimation and heat stress memory are inextricably linked. In this study, we performed a comprehensive RNA-Seq analysis on Arabidopsis seedlings that go through the heat acclimation, memory, heat shock and recovery in one time-course experiment. The results of this study present a systematic view of transcriptional changes, identifying genes and pathways that are implicated in heat acclimation and heat stress memory. We reveal that flavanols and genes involved in stomatal development are important elements of heat tolerance, which are targeted during the heat memory phase, leading to improved plant performance under heat stress.

## Results

### Heat acclimation increases survival rate after heat shock and the protective effect is time-dependent

To identify the most physiologically relevant protocol to study the dynamic responses of heat acclimation, we first examined the basal tolerance of 12 days old Arabidopsis seedlings (Col-0). The seedlings were exposed to acute heat shock (45°C) for 90 minutes (**Figure 1 B**). None of the plants survived after five days of recovery. The seedlings were also observed to die even when the heat shock treatment was reduced to 60 mins (**Figure 1 B**), suggesting the lethal nature of the abrupt increase in temperature. Interestingly when the seedlings were exposed to a gradual increase of temperature from 22°C to 45°C over a period of 6 h, followed by 90 minutes at 45°C, the survival rate was as high as for the plants grown under non-stress conditions (**Figure 1 A)**. The only visible difference between acclimated and non-acclimated plants was the slightly pale green coloring of the acclimated plants (**Figure 1 A**). The differences between non-acclimated and acclimated plants became more apparent when they were exposed to acute heat stress (**Figure 1 B**). Plants exposed to the acclimation treatment showed an increased survival rate compared to the non-acclimated plants. Those results show that gradual exposure to increased temperature results in acquired heat tolerance compared to the acute increase of temperature.

**Figure 1.**
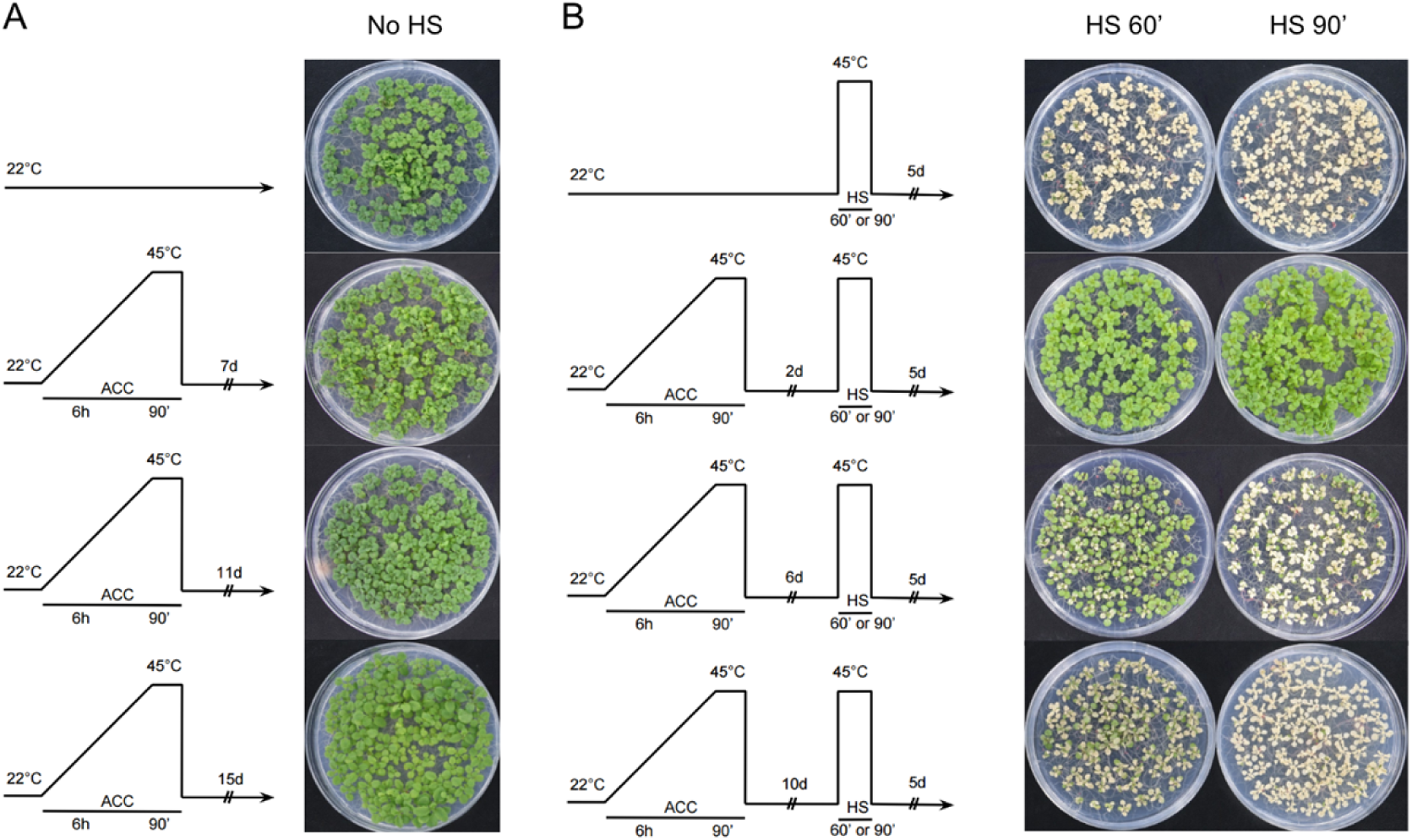
Different heat stress regimes to examine the effect of heat acclimation on Arabidopsis seedlings. **(A)** The temperature regime that plants were exposed to during the acclimation phase. The seedlings were exposed to acclimation (ACC) treatment 14, 10, or 6 days after germination and allowed to recover for different times (7, 11 and 15 days) before the pictures were taken. **(B)** The temperature regime that plants were exposed to during the acclimation phase and the heat shock period. The seedlings were exposed to acclimation (ACC) treatment 14, 10 or 6 days after germination, and allowed to recover for different times (2, 6 and 10 days) before heat stress was applied. The images were taken 5 days after the heat stress.

To examine how long this acquired thermotolerance (i.e. themomemory) can last, we subjected the acclimated plants to another abrupt heat stress changing the recovery period to 2, 6 and 10 days after the acclimation treatment. Acclimated plants showed the highest survival rate after exposure to the acute heat shock when the time gap between the acclimation and the heat shock was the shortest (2 days) (**Figure 1 B**). With increased length of the memory period after acclimation, the survival of the seedlings after acute heat stress decreased but was still higher than in non-acclimated plants (**Figure 1 B**). Those results indicate that the mechanisms involved in the acquired heat tolerance conferred by acclimation are time-dependent. The acquired thermotolerance was observed even after 10 days of the acclimation treatment (**Figure 1 B**). To our knowledge, 10 days is the longest thermomemory that has been observed to-date in Arabidopsis seedlings. These results demonstrate a clear impact of the exposure to gradually increasing stress level (acclimation) on increased tolerance to heat shock.

### Overview of transcriptome changes during the heat stress regime

To gain insights into the transcriptome-wide changes during heat acclimation and the subsequent heat stress response of acclimated and non-acclimated plants (T1 to T5), we sequenced the transcriptome of the acclimated plants during four time points of the acclimation phase, and one time point at two days (T5) into the memory phase. The transcriptome of the non-acclimated and acclimated plants was sequenced at the developmental stage corresponding to four days after acclimation, directly preceding the acute heat stress treatment (T6 and T9), 90 mins after application of heat stress 45°C (T7, and T10) and two days into the recovery phase after the heat stress application (T8 and T11) (**Figure 2 A**). Plant phenotypes of selected time-points can be found in **Supplementary Figure 1**.

**Figure 2.**
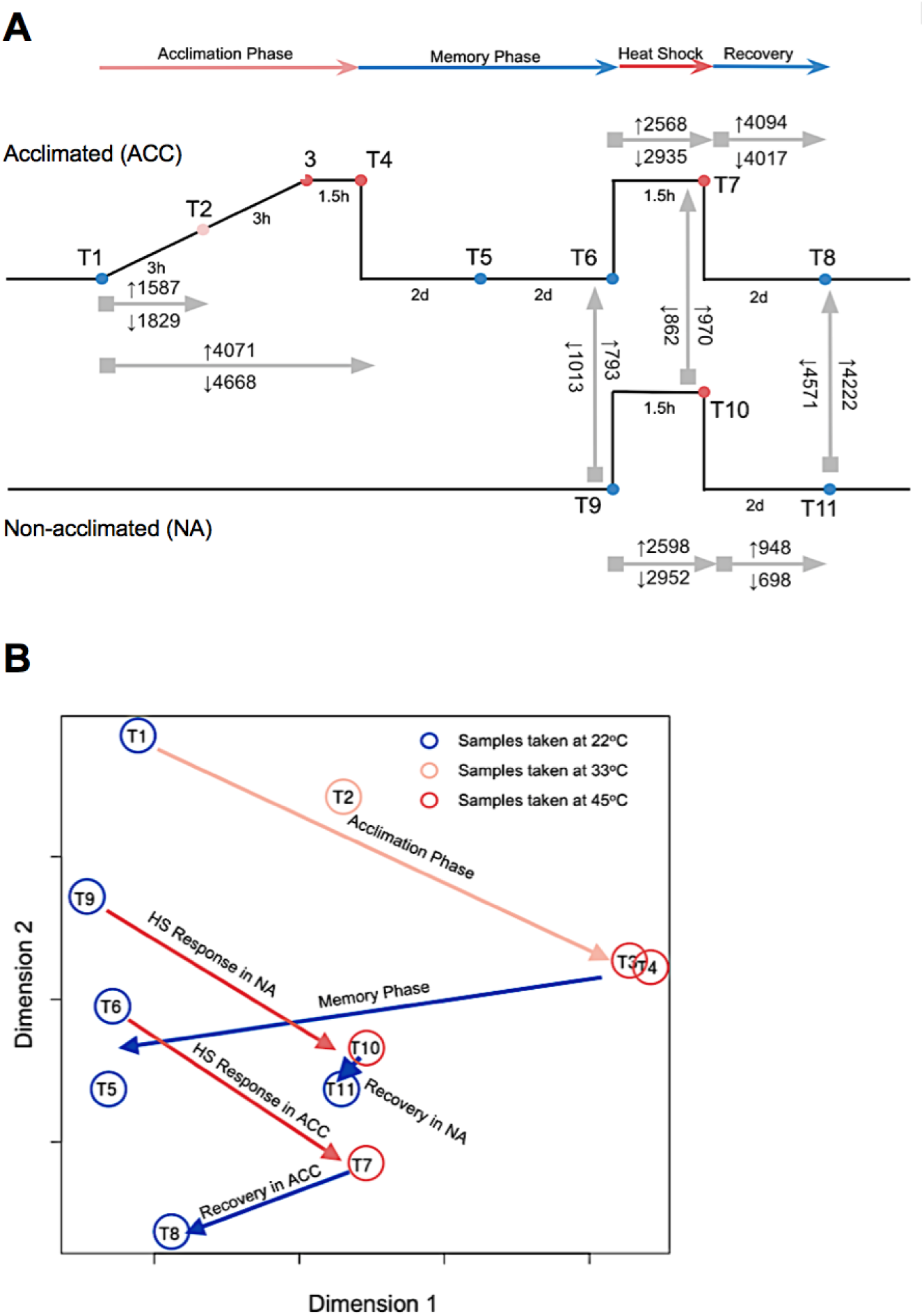
Overview of transcriptome changes during the heat stress regime. **(A)** Experimental design of RNA-Seq with the timepoints used for the RNA-Seq indicated with (T). The number of differentially expressed genes between selected time points in acclimated and non-acclimated samples are indicated above and below the arrows. **(B)** Overall expression characteristics of all time points visualized by multi-dimensional scaling plot. The distances between the points are approximately equal to their dissimilarities. Each time point represents the average expression of four biological replicates at a specific sampling point.

We selected pairwise time-points of plants undergoing acclimation as well as acclimated vs. non-acclimated plants during the heat shock and recovery phase, and identified the differentially expressed genes of each comparison (**Figure 2 A**). The number of differentially expressed genes (up and down from control with FDR p-value < 0.05) was calculated for each comparison (**Figure 2 A**). We also examined the similarities and differences in global changes in gene expression by applying a multi-dimensional scaling approach. The resulting two-dimensional map of relative distances between samples (**Figure 2 B**) revealed profound transcriptome reprogramming during the acclimation phase (T1 to T4), similar to the global patterns observed during the acute heat stress. During the memory phase (T5 and T6), the general expression profile was observed to revert to a pattern similar to the one observed in the non-acclimated plants (T6), but different from the original starting point (T1). This difference between acclimated (T6) and non-acclimated plants (T9) at similar developmental stage indicates transcriptional memory of the acquitted thermotolerance. During heat shock, regardless of the pre-treatment (acclimated vs. non-acclimated), the transcriptomes exhibited a similar response signature, reflecting similarities in general patterns in response to the heat shock (T7 or T10). Two days into the heat shock recovery, the acclimated plants showed a unique gene expression state (T8), that is more similar to that of the non-acclimated state (T5, T6), while the non-acclimated plants remained in the state that was closer to the pattern observed during heat stress treatment (T7 and T10).

### Transcriptional changes during the acclimation phase

The acclimation treatment induced gradual changes in the transcriptome, significantly altering the expression of 3416 and 8739 genes at 3 and 7.5 hours respectively into the acclimation treatment (**Figure 2 A**). The number of differentially expressed genes increased with the gradual increase in temperature, but the additional 90 minutes of heat stress does not result in additional changes to the general transcriptome profile (**Figure 2 B**).

Three hours after the onset of acclimation (T2), we identified 1587 genes that are up-regulated and 1829 of genes down-regulated compared to the starting non-heated state (T1), while at the end of the acclimation, the number of up and down differentially expressed genes are 4071 and 4668, accounting for 45% of the transcriptome (**Figure 2 A, Supplementary Table 2**). GO enrichment analysis indicated that the most enriched biological pathways from the up-regulated genes during the acclimation (T2 vs T1, and T4 vs T1) are the categories “response to heat”, “response to high light intensity”, “protein folding”, “response to hydrogen peroxide”, “response to oxidative stress”, “response to reactive oxygen species” (**Supplementary Table 2**). HS-responsive genes are commonly shared in those ontologies, such as various classes of heat shock proteins and antioxidant enzymes. Biological processes of “response to stimulus”, “phosphorylation”, “cell division” and “developmental process” were enriched in the transcripts repressed during the acclimation (**Supplementary Table 3**).

Visualization of transcriptional change during the acclimation phase using MAPMAN (Thimm et al., 2004) revealed biological processes related to heat acclimation (**Figure 3**). Consistent with the GO enrichment results, we observed significant up-regulation of abiotic stress related genes during acclimation period (**Figure 3 A**), and a large proportion of genes are down-regulated in cell division, cell cycle and developmental process at the end of acclimation (**Figure 3 B**). Those results suggest that exposure to a gradual increase in temperature slows plant development. Interestingly, we also observed profound repression in genes involved in biotic stress response at the end of acclimation (**Figure 3 B**), pointing to a tradeoff between biotic resistance and thermotolerance acquisition.

**Figure 3.**
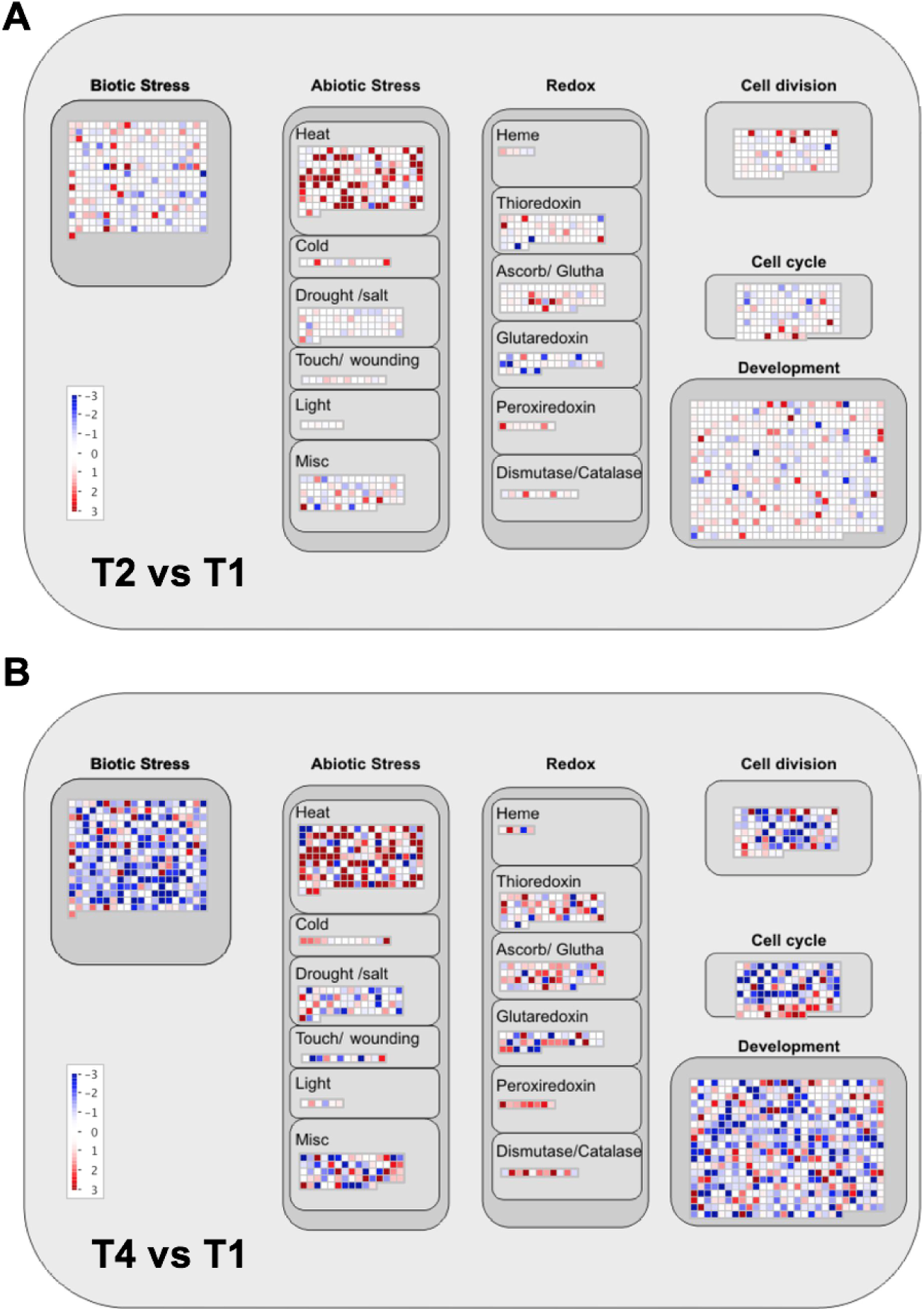
MAPMAN visualization of the transcriptional changes during the acclimation phase. The comparison of mapped 1597 gene expression in the MAPMAN cellular response pathway between the start of the acclimation and **(A)** 3h into the heat acclimation (T2 vs T1) and **(B)** at the end of the acclimation (T4 vs T1). Each square represents one gene mapped to a functional category in the cellular response pathway. Blue color: up-regulated; Red color: down-regulated.

### Transcriptional changes underlying heat stress memory

During the memory phase (T5 to T6), the transcriptional pattern of the acclimated plants gradually reverts to the state that is globally similar to the non-acclimated state (T9) (**Figure 2 B**). However, the general transcriptome is still clearly different with about 1800 differentially expressed genes between the acclimated and non-acclimated plants at the same developmental stage, before the heat shock (**Figure 2 A**).

Intriguingly, among the 793 up and 1089 down-regulated genes identified to be differentially regulated between acclimated and non-acclimated plants at the end of the memory phase (T6 vs T9), only about one-third of them (254 in the up-regulated, and 367 in the down-regulated) are shared with the genes which showed altered expression in response to acclimation (T1 vs T4), implying a distinct transcriptional program during the memory phase (**Figure 4 A**). To visualize the biological processes that are significantly affected at the end of the memory phase, we constructed GO enrichment map (**Figure 4 B**) which shows the functional categories enriched in the acclimated plants compared to the non-acclimated seedling at the same developmental stage (T6 vs T9). Enriched GO categories in the up-regulated genes include “response to stimulus”, “flavonoid biosynthetic process”, “response to heat”, and “lipid transport”, while the down-regulated genes are enriched in “photosynthesis”, “response to abiotic stimulus”, and general metabolism (**Supplementary Table 4, 5**). We were particularly intrigued by the significant overrepresentation of genes associated with the flavonoid biosynthetic process in the acclimated plants at the memory phase (T6 vs T9) (**Figure 4**), as flavonoids were previously not reported to have a role in heat tolerance. Furthermore, genes in this pathway are significantly over-represented in the memory phase. Therefore, we decided to examine the contribution of the flavonoid pathway in thermotolerance.

**Figure 4.**
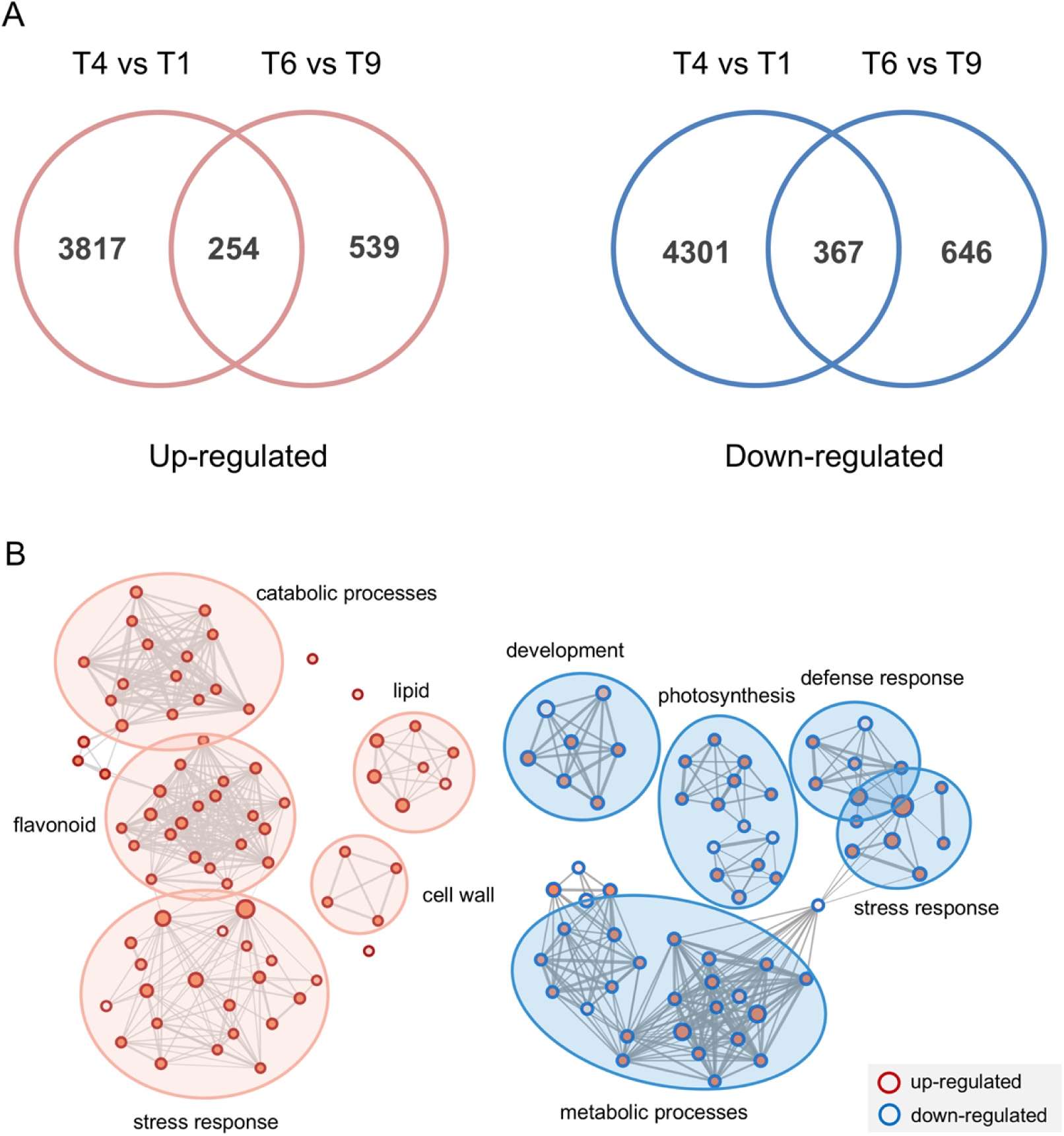
Transcriptional changes underlying heat stress memory. **(A)** The number of genes up and down-regulated at the end of acclimation (T4 vs T1) and at the end of the memory phase in acclimated plants relative to the non-acclimated plants (T6 vs T9). **(B)** GO enrichment map of genes up and down-regulated in acclimated plants four days after acclimation relative to the non-acclimated plants. The analysis was performed in Cytoscape using BiNGO and Enrichment Map plugin. GO categories that were significantly overrepresented among the differentially expressed genes between T6 and T9. Each node represents a biological pathway in the GO category. The different grades of red of the circles correspond to the level of significance of the overrepresented GO category. The size of the circles is proportional to the number of genes in each category. Edge (gray line) size corresponds to the number of genes that overlap between two GO categories.

### Flavonoid and its role in heat stress

The expression of genes encoding for enzymes of the central flavonoid pathway in acclimated and non-acclimated plants was examined in more detail (**Figure 5**). We observed that among all the key flavonoid biosynthesis genes, the transcript abundance was higher in acclimated than in non-acclimated plants. The same trends were sustained during the heat shock, and two days after returning to ambient temperature. Moreover, two main regulators of the flavonoid biosynthesis pathway in Arabidopsis, production of anthocyanin pigment 1 and 2 of the MYB transcription factor family (PAP1, AT1G56650; PAP2, AT1G66390) (Tohge et al., 2005; Ilk et al., 2015), were also found to be up-regulated in acclimated plants. A thorough investigation at all time-points shows the induction of PAP1 and PAP2 occurred during acclimation and increased even further two days into the memory phase (T5) (**Supplementary Figure 4**). Similar increases in transcript abundance during the memory phase (T5 vs T4) were also found in flavonoid synthetic *Transparent Testa* (*TT*) genes *TT4, TT5, TT6* and *TT7* (**Supplementary Figure 4**).

**Figure 5.**
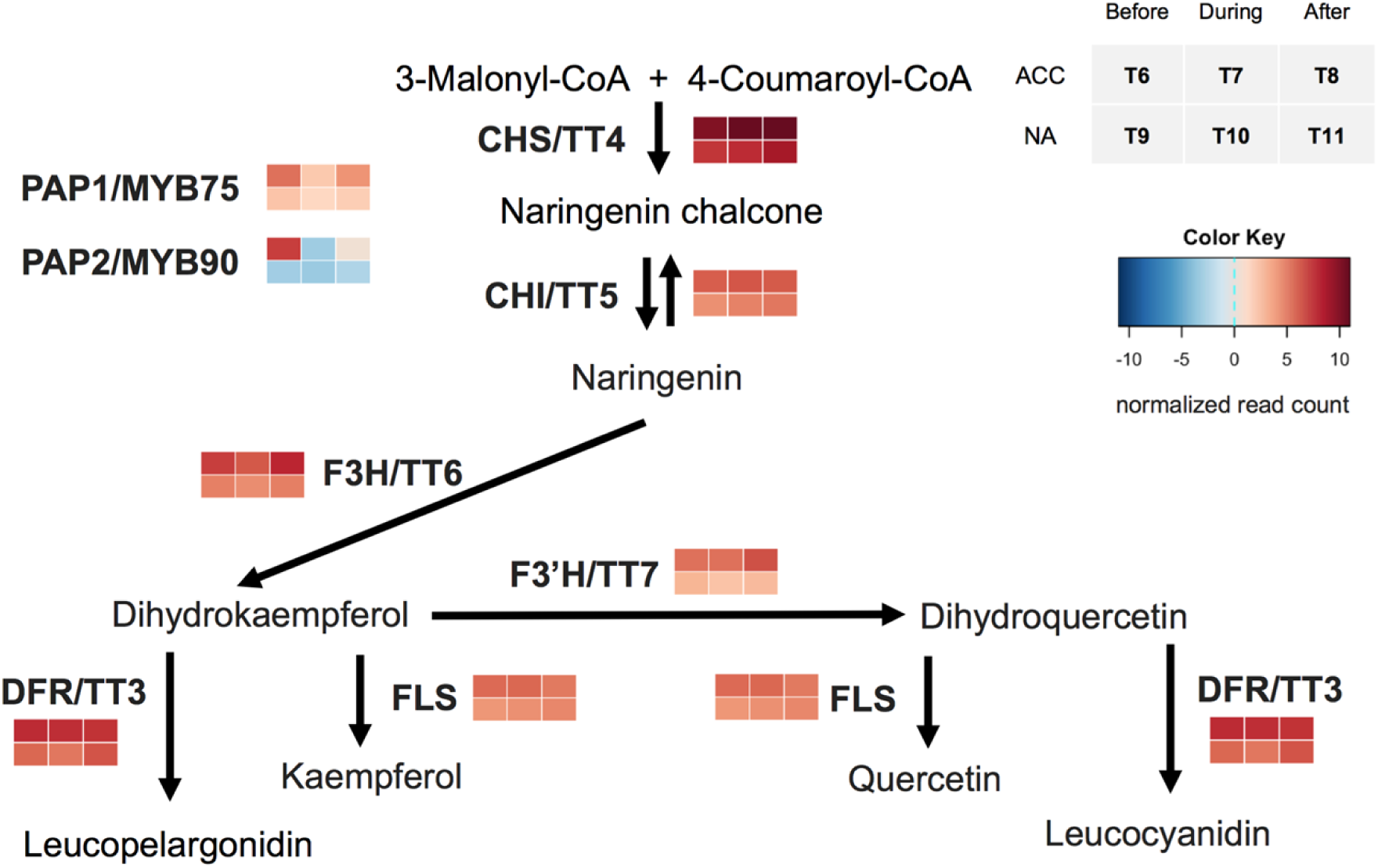
Expressions of key flavonoid biosynthetic genes before, during and after the heat shock in acclimated and non-acclimated plants. Color key represents the average normalized read count in log2 from four biological replicates. Genes encoding central enzymes of the flavonoid pathway are: AT5G13930 CHS/TT4, chalcone synthase; AT3G55120 CHI/TT5, chalcone isomerase; AT5G42800 DFR/TT3, dihydroflavonol reductase; AT3G51240 F3H/TT6, flavonol 3-hydroxylase; AT5G08640 FLS, flavonol synthase; TT, transparent testa. Genes encoding regulators of flavonoid pathway are: AT1G56650 PAP1/MYB75 and AT1G66390 PAP2/MYB90.

Several conceivable mechanisms that might explain the role flavonoids might have in plant stress tolerance. The scavenging of reactive oxygen species (ROS) is one of them. It has been shown that flavanols, especially quercetin, are potent antioxidants (Rice-Evans et al., 1997), and supplying quercetin externally to plants can protect them from oxidative damage (Kurepa et al., 2016). To test whether quercetin accumulation confers better thermotolerance in Arabidopsis, we tested the basal thermotolerance on 7 days old Col-0 seedlings grown on ½ MS media with or without supplementation of 200μM quercetin. By examining the survival four days after heat shock treatment, we observed that quercetin improved the seedlings’ survival after heat shock (**Figure 6 A, B**). Additionally, we screened the basal thermotolerance of several *tt* mutants (*tt4*, SALK_020583; *tt5*, SALK_034145; *fls*, SALK_106244C1), which were previously described to accumulate less quercetin (Bowerman et al., 2012). While *tt3* and *fls* exhibited similar survival rates to WT seedlings (**Figure 6 C, D**), *tt4* seedlings were showing higher basal thermotolerance than the WT. These results indicate that flavanols might have a role in heat shock and contribute to acquired thermotolerance.

**Figure 6.**
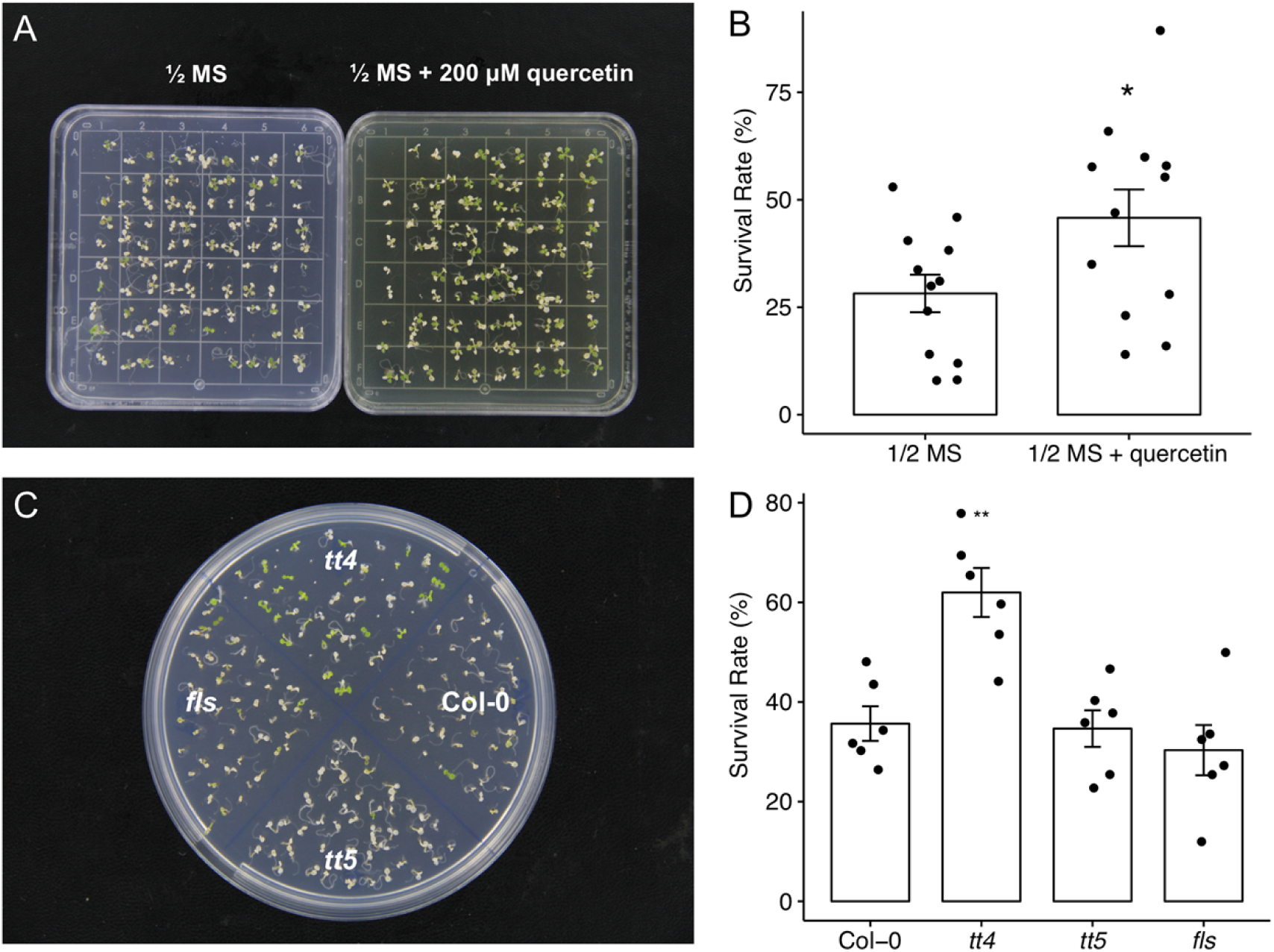
Quercetin affects basal thermotolerance of Arabidopsis seedlings. **(A)** 100 seedlings of Col-0 were germinated on media containing ½ MS with or without the addition of 200 uM quercetin for 7 days. The survival of the seedlings after exposure to 45°C for 45 minutes was determined 3 days after the plants were returned to the ambient growth chamber for recovery. **(B)** The seedling survival rate was established based on the survival rate scored over twelve plates divided over three experiments. **(C)** Col-0 and *tt* knockout mutants were germinated on ½ MS media for 7 days and subjected to 45°C for 45 min. The survival of the seedlings after exposure to 45°C for 45 minutes was determined 3 days after the plants were returned to the ambient growth chamber for recovery. **(D)** The seedling survival rate was established based on the survival rate scored over six plates divided over two experiments. The bars represent means, while the error bars represent SE. Significant differences, evaluated by student t-test, are marked by asterisks: *p < 0.05, **p < 0.01.

### Heat acclimation affects cooling capacity during the recovery and results in reduced water loss in acclimated plants

One of the most important processes that we hypothesized would be in operation during acclimation is the ability of the plants to acquire a more efficient cooling mechanism. Therefore, we visualized the leaf surface temperature of acclimated and non-acclimated plants using thermo-imaging after the heat treatment. Both individual leaf and the whole rosette collected from acclimated plants showed lower leaf surface temperatures than non-acclimated ones during the cooling period, indicating improved cooling capacity (**Figure 7 A**). The above observations imply heat acclimation affects the leaf cooling capacity, suggesting differences in stomatal development and/or functionality.

**Figure 7.**
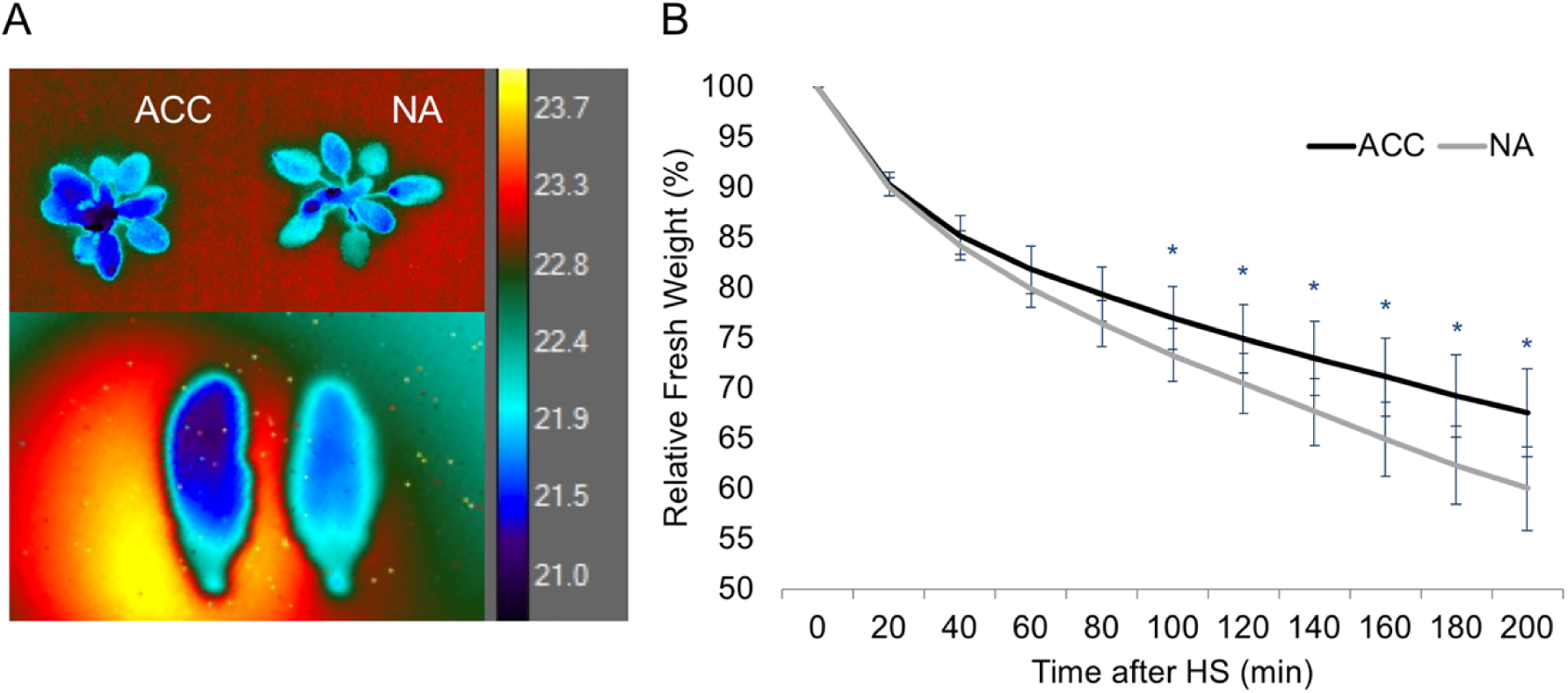
Relative fresh weight and leaf temperature of acclimated (ACC) and non-acclimated (NA) plants after the heat treatment. 4-week-old soil-grown plants were treated with or without heat acclimation. Four days later, they were subjected to heat stress (45°C for 90 mins). **(A)** Surface temperatures of the rosette or individual leaf during recovery from heat stress were recorded using a thermal camera (FLIR A655SC). Left: acclimated plant; Right: non-acclimated plants. The color key of the temperatures was indicated on the right. **(B)** Their aerial parts were excised and weighed for their fresh weights every 20 mins after the heat stress at 22°C under 65% relative humidity environment. Data points represent means ± SE of four biological replicates. *indicates significant differences between acclimated and naïve plants with p-value < 0.05 as determined by Student’s t-test.

To gain insights into the physiological changes heat acclimation induced to Arabidopsis, the differences in the transpiration rates were examined between acclimated and non-acclimated mature plants. Four-week-old plants grown in soil were exposed to the acclimation treatment. Four days after, acclimated and non-acclimated plants were exposed to 90 minutes of heat shock (45°C) and the fresh weight of the whole Arabidopsis rosette was measured every 20 minutes over 200 minutes. We observed that the loss of relative fresh weight was slower in the acclimated plants than in the non-acclimated plants (**Figure 7 B**). The results suggest that acclimation treatment leads to physiological changes in the leaves that help water loss prevention upon recurring heat.

### Heat acclimation affects stomatal development and function

The reduced rate of transpiration in acclimated plants could result from differences in stomatal size and/or stomatal aperture. To test whether heat acclimation affects stomatal development and function, we measured the stomatal density and stomatal aperture of acclimated and non-acclimated plants four days after the heat acclimation. We observed that acclimated plants showed a reduced number of stomata and smaller stomatal apertures compared to non-acclimated plants (**Figure 8 A, C**). These results are in line with the reduced transpiration rate we observed previously in water loss assay (**Figure 7 A)**, and suggest that heat acclimation leads to more efficient plant water retention.

**Figure 8.**
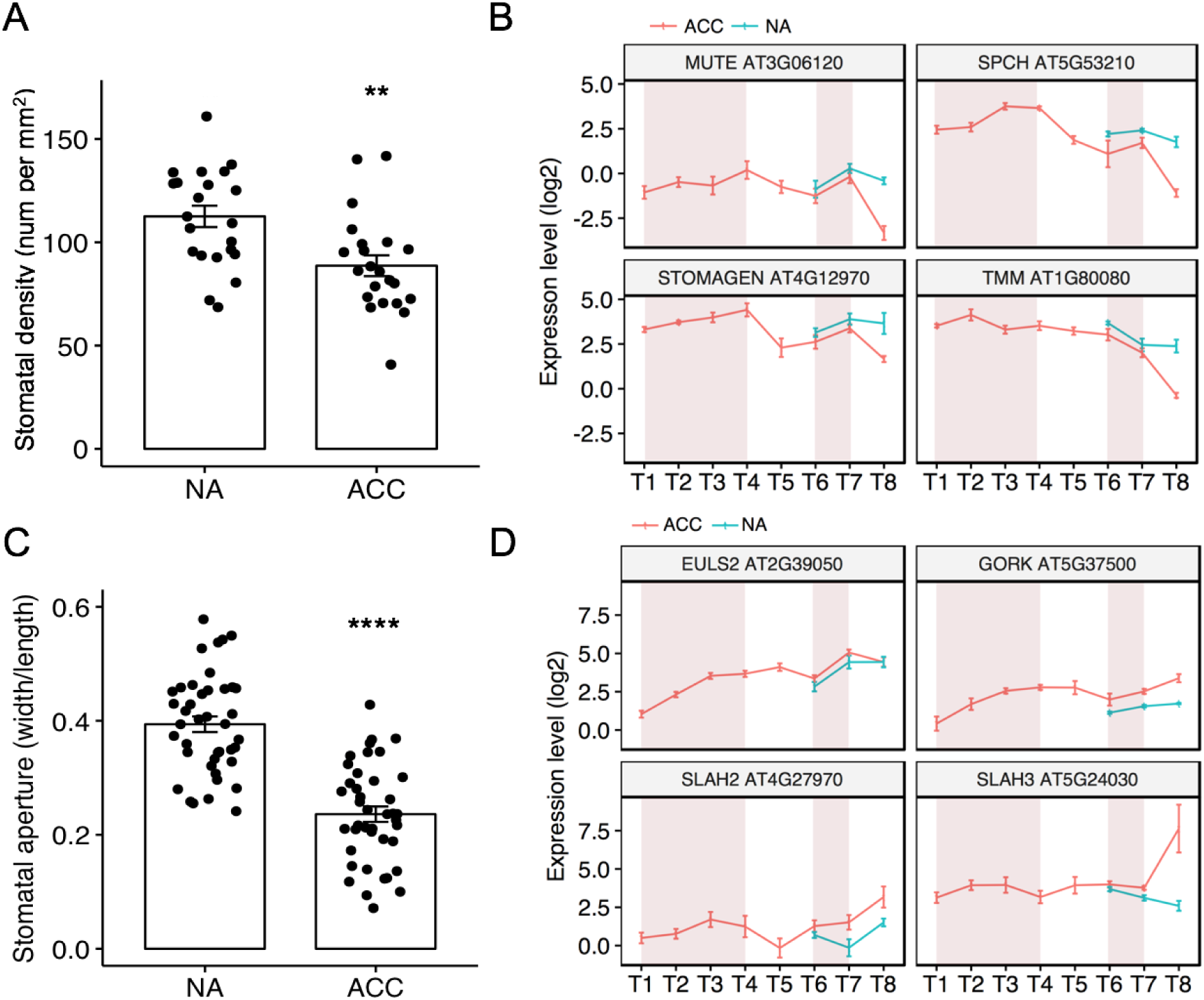
Heat acclimation affects stomatal development and function. 4-week-old soil-grown Col-0 plants were exposed to heat acclimation. **(A)** Average stomatal density and (**C**) stomatal aperture was measured from the adaxial surface of the fifth rosette leaves were determined four days after the acclimation treatment. The bar graphs represent mean, and error bars represent SD of image counts from four individual plants per condition. Significance levels are calculated by student’s t-test: *p<0.05, **p<0.025, ***p<0.01, and ****p<0.005. Average expression of normalized read counts of key **(B)** stomatal development genes and **(D)** functional regulators of stomatal aperture. Data points represent the mean ± SE of four biological replicates. Red line corresponds to the expression pattern observed in acclimated plants. Blue line represents the expression pattern observed in non-acclimated plants. Pink background indicates the acclimation period (T1 to T4) and the heat shock period (T6 to T7).

We subsequently examined the transcriptional patterns of genes known to be involved in stomatal development in our RNA-Seq dataset during the heat acclimation and heat stress regime. Three bHLH family transcription factors (MUTE, SPEECHLESS and STOMAGEN), previously described to promote the initiation, proliferation and terminal differentiation of cells in the stomatal lineage (MacAlister et al., 2007), were observed to be repressed after the heat acclimation (T4 to T6), and their transcripts were lower than in non-acclimated plants for the rest of the heat regime (T6 to T8) (**Figure 8 B**). The expression of TMM, which promotes the division of stomata cells (Nadeau and Sack, 2002) showed a similar pattern to the bHLH genes. Interestingly, during the heat shock recovery period (T7 to T8), there is an even sharper decline of expression of the four genes in the acclimated plants, while the non-acclimated plants do not respond to the same degree (**Figure 8 B**).

In addition, an examination of the expression of genes involved in regulation of stomatal aperture, including outward rectifying potassium GORK channel in the guard cells (Hosy et al., 2003; Schroeder, 2003), the EULS3 lectin (Van Hove et al., 2015), and the SLAH anion channel (Geiger et al., 2011) was undertaken. We observed an acclimation induced up-regulation of the transcript abundance of genes contributing to stomatal closure (**Figure 8 D**). These transcripts remained more abundant in acclimated plants than non-acclimated ones at the later time points (T6 to T8).

In summary, the decline of transcript abundance of the main stomatal regulators might serve as a signal to decrease stomatal development as an adaptive strategy for heat. The up-regulation of stomatal aperture regulators after heat acclimation suggests that increased efficiency of stomatal aperture control might be an acquired and adaptive mechanism, enabling plants to cope with elevated temperatures.

### *AGL16* is responsive to heat and regulates thermotolerance

To test the role of stomata development in acquired thermotolerance we focused on the gene reported to be a master regulator of stomatal development, MADS-box transcription factor AGAMOUS-LIKE16 (AGL16). AGL16 is found to be highly expressed in guard cells and trichomes in Arabidopsis (Alvarez-Buylla et al., 2008) and is targeted by miR824 in the regulation of satellite meristemoid lineage of stomatal development (Kutter et al., 2007). We found that the transcripts of primary miR824 were up-regulated during heat acclimation and heat shock treatment. The heat stress itself also resulted in the up-regulation of pri-miR824 in both acclimated and non-acclimated plants (**Figure 9 A**), suggesting that the ability to upregulate the miR824 was not restricted to acclimated plants. For *AGL16*, we noticed a slight down-regulation in the first 3h of the acclimation (T2 to T1), and two days after the acclimation (T5 to T4) (**Figure 9 C**). The expression of *AGL16* remained repressed in acclimated plants compared to the non-acclimated plants. Both acclimated and non-acclimated plants showed a clear reduction of *AGL16* expression during the heat shock treatment, consistent with the up-regulation of *pri-miR824*. This heat induced up-regulation of miR824 and down-regulation of AGL16 was further confirmed by qRT-PCR, by subjecting one-week-old seedlings directly to abrupt heat stress (**Figure 9 B, D**). These results suggest that the repression of *AGL16* occurs at both acclimated and non-acclimated plants, and possibly contributes to the differences in stomatal development which results in acquired heat tolerance. When the *agl16* seedlings were tested for basal thermotolerance, they exhibited a higher survival rate than WT, confirming that the acclimation-induced downregulation of AGL16 promotes the acquisition of thermotolerance (**Figure 9 E, F)**. The data therefore, point to a novel mechanism of acquired thermotolerance, where the increased temperature leads to a transcript accumulation of *pri-miR824*, repressing AGL16, resulting in reduced stomatal development and hence improved survival in the event of recurring heat stress.

**Figure 9.**
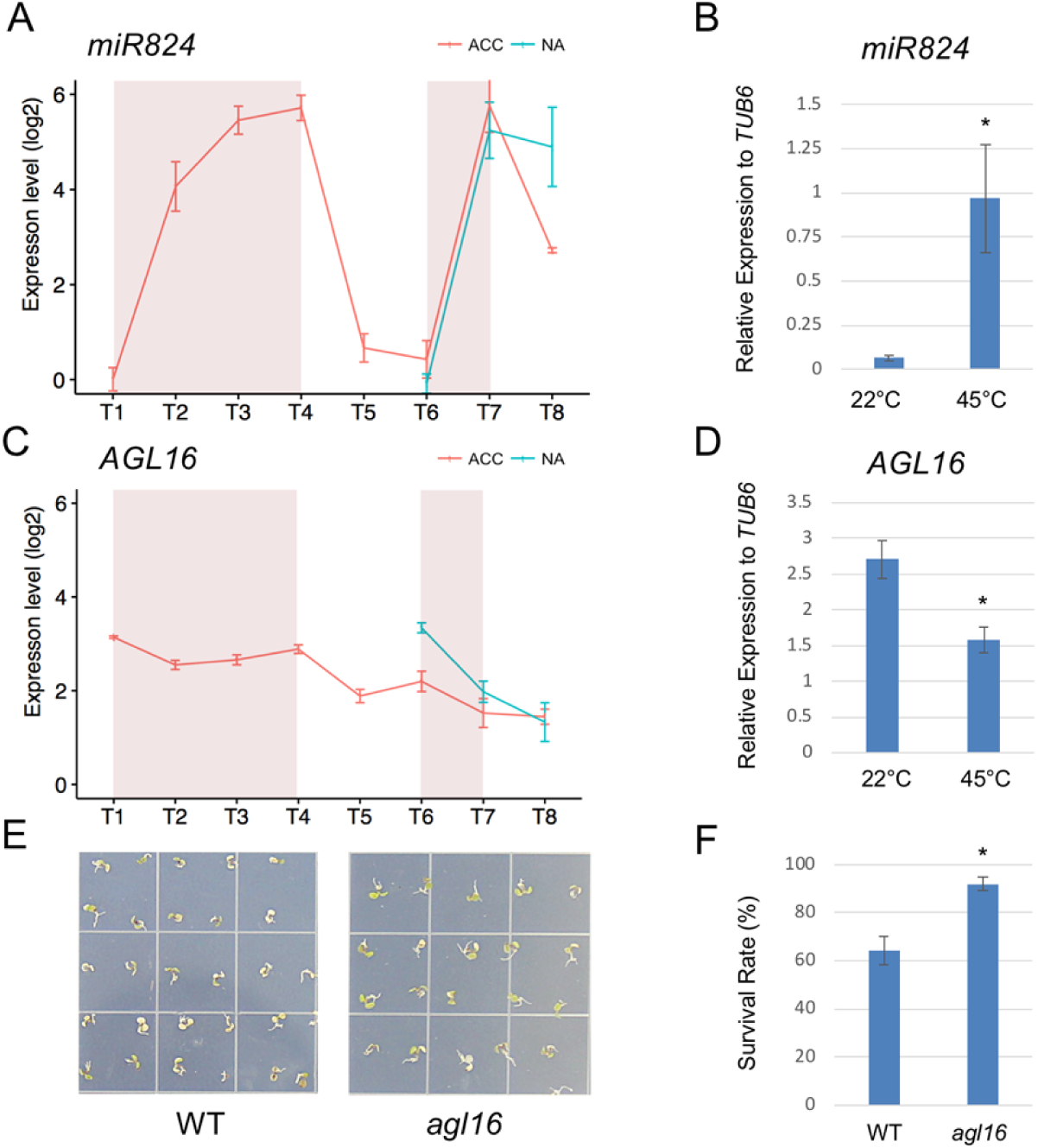
The repression of *AGL16* expression under heat results in enhanced thermotolerance. Expression levels of *AGL16* (**A**) and *miR824* (**C**) of acclimated and non-acclimated plants of all time-points during our RNA-Seq experiment were shown as normalized counts per million in log2 scale. Each data point corresponds to the mean and standard error of four biological replicates. ACC: acclimated plants. NA: non-acclimated plants. Pink background indicates the acclimation period (T1 to T4) and the heat shock period (T6 to T7). (**B**) The real-time PCR analysis of *AGL16* and *pri-miR824* expression in the wild-type before the heat treatment (22°C) and 60 mins after the heat treatment (45°C). The expression level of housekeeping gene TUB6 was used as the reference. Each data point corresponds to the mean and standard error of three biological replicates. (**D-E**) Seven-day-old seedlings grown on ½ MS plates at 22°C were shifted to 45°C for 60 min and then returned to 22°C. The plants were photographed four days after the heat treatment and survival rates of wild-type (WT) and *agl16* seedlings after heat shock were scored. Each value is the mean and standard error of three biological replicates (n =100 seedlings per replicate).

## Discussion

Temperature is fluctuating in natural conditions. Plants often experience temperatures above their optimal growth condition, which requires acclimation to altered conditions in a relatively short time. Previous studies have shown that a gradual increase of temperature elicited acquired thermotolerance than a step-wise increase of temperature, and this might reflect a more general mechanism that contributes to the homeostasis of metabolism and heat adaptation of plants. (Larkindale and Vierling, 2008; Mittler et al., 2012). The experiments shown in this study show that this acquired thermotolerance can sustain for up to 10 days, but the strength of the protective mechanism, so-called ‘stress memory’, weakens with the increased length of the time interval between acclimation and repeated stress exposure (**Figure 1**).

As the heat stress memory is transient and time-dependent, we hypothesized that a large portion of the acquired thermotolerance is due to transient transcriptional changes. General patterns that we observed during the acclimation (**Figure 2**) pinpointed that the transcriptome reprogramming happens very fast during the gradual increase of temperature. Although our experimental design does not allow us to distinguish between the transcriptional changes caused by circadian rhythm during the acclimation process, it nevertheless allowed us to gain insight into general patterns and enrichment of transcriptional changes during acclimation. The most prominent transcriptional responses during the heat acclimation included the activation of heat shock transcription factors and heat shock proteins, which assist protein homeostasis through their chaperone activities. There are 21 HSFs in *Arabidopsis thaliana*, so far eight have been shown to act in the responses to heat (Charng et al., 2007; Schramm et al., 2007; Ikeda et al., 2011; Liu et al., 2011; Scharf et al., 2012). We have detected 17 of them in our dataset (**Supplementary Figure 3**). Six of them (*HSFA2, HSFA7A, HSFA7B, HSFB2a, HSFB2b, and HSFB4*) are responding quickly during the onset of temperature increase (T2 vs T1). Notably, *HSFA2*, the master regulator of heat response and formation of acquired thermotolerance (Charng et al., 2007; Lämke et al., 2016) shows great induction in expression during the acclimation, but it also activated to the similar level both in non-acclimated and acclimated plants upon the recurring heat shock treatment (**Supplementary Figure 3**). The expression of *HSFA1e* was observed to be up-regulated during the acclimation phase (T2 to T4) and was induced to a much higher level during the reoccurring heat stress exposure in acclimated plants than in the non-acclimated plants (**Supplementary Figure 3**). The exact role and mechanisms of HSFA1e in thermomemory remains to be investigated. We hypothesize that the more pronounced induction of *HSFA1e* during reoccurring heat stress contributes to the thermotolerance, possibly assisted by epigenetic markers and the activation of the classical HSFA2-initiated heat stress response. Most of the transcriptional changes in response to moderate levels of heat stress are transient and restored when the environmental stimulus is removed (Larkindale and Vierling, 2008; Stief et al., 2014). We observed a number of genes for which the transcripts were more abundant during the acclimation and remained high throughout the memory phase. These memory patterns were exhibited by the transcripts of APX2, FTSH6, HSP21, HSA32 and MBF1C (**Supplementary Figure 5**), supporting their roles in thermomemory that have been described previously (Charng et al., 2006; Suzuki et al., 2008; Sedaghatmehr et al., 2016).

The transcriptional state during the memory phase changed to something similar to that of non-acclimated plant, but yet distinct. At the end of the memory phase, the up-regulated transcripts were enriched for the GO terms related to the flavonoid pathway. A recent metabolome study has identified the significant accumulation of two flavonoids (dihydrokaempferol and naringenin) after heat acclimation (Serrano et al., 2019). The positive correlation of flavonoid gene expression and metabolite has also been established in cold acclimated Arabidopsis (Hannah et al., 2006; Schulz et al., 2016). Flavonoids are known to have an important role in the response to freezing (Korn et al., 2008; Schulz et al., 2016) and UV-B tolerance (Falcone Ferreyra et al., 2012; Emiliani et al., 2013), but their roles in heat tolerance was not explored so far. Flavonoids exhibit a great potential to inhibit the generation of ROS, acting as ROS scavengers (Agati et al., 2012; Nakabayashi et al., 2014) and also have direct effects on the stability of cellular membranes (Movileanu et al., 2000; Pawlikowska-Pawlęga et al., 2014). One of the most abundant flavonoids in plants, quercetin, was found to affect membrane fluidity, cooperativity and the temperature of phase transition (Tsuchiya et al., 2002; Pawlikowska-Pawlęga et al., 2007). Quercetin treated seedlings also exhibited higher survival rates in response to heat stress treatment (**Figure 6 A**), possibly through enhanced anti-oxidation activity and/or changes in membrane fluidity. Further experiments will be necessary to evaluate the quercetin contribution to ROS scavenging and/or and maintenance of cell membrane integrity during heat stress. While the majority of the transcripts of flavanol biosynthetic components exhibited higher abundance in acclimated seedlings (**Figure 5**), two of the three tested flavanol mutants did not show a difference in heat tolerance (**Figure 6 C, D**). The lack of significant differences in flavonoid biosynthesis mutants used in this study could be explained by the complex character of thermotolerance and the functional redundancy of flavonoid-related proteins and metabolites. The fact that only the *tt4* mutant showed enhanced heat stress tolerance (**Figure 6 C, D**) somehow seems to contradict the results obtained by quercetin supplementation (**Figure 6 A, B**) and the transcriptional profiles. This could be due to an altered accumulation of other flavonoids with potential compensatory effects. The contribution of a single gene or pathway to the total phenotype is therefore very difficult to quantify. Future studies quantifying changes in the flavonoid compounds during the acclimation and heat stress, accompanied by the physiological studies of multiple knock-out mutants of several flavonoid biosynthetic components might provide a more comprehensive understanding of the role of flavanol in acquired thermotolerance.

Efficient cooling is an essential mechanism, necessary for the plant to survive at high temperatures. The known mechanisms of cooling occur through the evaporation through stomata. Plants can adapt to changing environmental conditions by adjusting the density and/or the aperture of the stomatal pores. Arabidopsis plants grown at higher temperature were previously observed to develop leaves with lower stomatal density, yet higher evaporative leaf cooling capacity (Crawford et al., 2012). However, the short-term thermal acclimation was not previously observed to induce adaptive developmental changes in stomatal density. Here, we observed that the transcripts of stomatal development genes were repressed after short-term heat acclimation, resulting in a measurable decrease of stomatal density within 4 days in heat-acclimated Arabidopsis plants (**Figure 8**), with decreased water loss and yet enhanced leaf cooling capacity (**Figure 7**). In addition, our thermal assay experiment with the *agl16* mutant confirmed that AGL16 is part of the acquisition of thermotolerance (**Figure 9 E, F**), most probably by affecting stomatal development. The AGL16 itself is tightly-controlled by miR824 (de Meaux et al., 2008; Hu et al., 2014), which showed dynamic induction in response to increased temperature in both acclimation and heat stress treatment (**Figure 9 C, D**). Collectively, our results suggest that stomata development and regulation of aperture are important targets for acquired thermotolerance in Arabidopsis.

In summary, in this study, we present a robust acclimation protocol and establish transcriptional signatures during acclimation, the memory phase, and subsequent heat shock. We confirmed some known processes underlying acquired thermotolerance and suggested functional roles for flavonoid biosynthesis and stomatal regulation in acquired thermotolerance. Our work provides a useful framework for future studies of acquired thermotolerance and a resource for future investigation of molecular mechanisms that govern heat acclimation and heat stress memory.

## Material & Methods

### Plant material

*Arabidopsis thaliana* seeds were surface-sterilized using 70% ethanol for three mins and washed with sterile water three times. 80 to 90 sterile seeds were evenly sowed on round 100 × 15 mm plastic petri dishes. Each plate was filled with 50 ml autoclaved media containing 0.22% (w/v) of MS basal salts, 1%(w/v) sucrose, 1% (w/v) Agar, with pH adjusted to 5.8 with KOH. The seeds were incubated at 4°C in dark for 3 days to ensure synchronized germination and transferred to the growth chamber (Percival) with 22°C under 16h / 8h of light / dark cycle (∼100 μmol m−2 s−1) with 60% relative humidity. T-DNA insertion lines of TT4 (AT5G13930), TT5 (AT3G55120), FLS (AT5G08640), and AGL16 (AT3G57230) are *tt4* SALK_020583; *tt5* SALK_034145; *fls*, SALK_106244C1; *agl16,* SALK_104701C respectively. They were ordered from the Nottingham Arabidopsis Stock Center (NASC).

### Heat acclimation and heat shock application

The acclimation treatment was applied by subjecting plates with Arabidopsis seedlings into the growth chamber (Percival) with the temperature gradually rising from 22°C to 45°C over the course of 6 hours (starting at the beginning of the photoperiod) in the light and keep at 45°C for 90 mins. Subsequently, the plates were transferred back to a growth chamber with standard (22°C) temperature. For the heat shock application, the growth chamber (Percival) was pre-heated to 45°C with lighting (∼100 μmol m−2 s−1) and 60% relative humidity. Each plate containing 80 acclimated or non-acclimated seedlings was transferred from the control growth condition (22°C) chamber to a heated growth chamber (45°C) for 90 mins from 1:30 pm to 3:00 pm (7.5 - 9h after the beginning of the photoperiod).

### RNA-Seq sampling, library preparation, and sequencing

We collected samples of 11 time-points from two batches of experiments. For each batch, 11 plates each containing about 80 germinated Arabidopsis seedlings were divided into the acclimated and non-acclimated group. At each sampling time-point from the experimental design (Figure 2.2 A), all leaf tissue from the designated plate was harvested and divided into two separate Eppendorf tubes, snap-frozen in liquid nitrogen and stored at -80°C. At day 12 after seed germination, T1 samples were taken right after the start of the photoperiod and before acclimation treatment starts. Plates from acclimated group were exposed to the heat acclimation treatment and samples were taken after 3 h (T2), 6 h (T3) and 7.5 h (T4), while other plates were returned to normal growth conditions and samples were collected two days (T5) and four days (T6) after at 1:30 pm. At day 16, the remaining acclimated and all the non-acclimated plants were exposed to heat shock treatment (45°C for 90 min), and samples were taken before the heat treatment (T7 for acclimated seedlings, T9 for non-acclimated seedlings), at the end of the heat shock (T8 for acclimated seedlings, T10 for non-acclimated seedlings) and two days after the application of heat stress at 1:30pm (T9 for the acclimated seedlings, T11 for non-acclimated seedlings). In total, we collected 44 samples from two independent experimental batches; with four biological replicates for each time point. RNA extraction was performed for each sample using RNeasy Plant Mini Kit (Qiagen). The RNA concentration was measured by NanoDrop 2000 spectrophotometer (ND-2000; Thermo Fisher Scientific). RNA integrity was assessed on an Agilent 2100 Bioanalyzer (Agilent Technologies). 2μg of RNA for each sample was used to construct mRNA library following the protocol of Illumina’s TruSeq Stranded mRNA Library Prep Kit. One sample from T3 was missing, due to failure in library construction. In total, 59 Gb paired-end reads of 21 samples from batch one, and 22 samples of 30 Gb single-end reads from batch two were generated (**Supplementary Table 2.1**).

### Processing of RNA-seq reads

Tophat (Kim et al., 2013) was used to map the raw RNA-seq reads onto the *Arabidopsis thaliana* reference genome (TAIR10), and htseq (Anders et al., 2015) was applied to quantify the transcript level for each gene. Hierarchical clustering analysis was performed to test for relative relatedness of all 43 samples from two experimental batches (**Supplementary Figure 2.2**).

### Identification of differentially expressed genes

Read counts mapped to individual gene models were calculated into cpm (counts per million), and genes with low read counts (cpm < 1 in more than two samples) were removed. Voom (Law et al., 2014) function from linear models for microarray and RNA-Seq data (LIMMA) framework was applied to convert raw read counts to log_2_(cpm) values with associated weights. Read counts were modeled with a negative binomial distribution, the common dispersion parameter was estimated using Empirical Bayes techniques. Pairwise comparisons between groups were made by using a quantile-adjusted conditional maximum likelihood (qCML) estimate. DecideTest from the LIMMA framework was used to test for differentially expressed genes (with adjusted p-value < 0.05).

### Functional enrichment analysis

AgriGO (Tian et al., 2017) was applied to determine which Gene Ontology (GO) categories were statistically overrepresented in biological pathway, cellular component and molecular function for each group of differentially expressed genes. All annotated genes in the Arabidopsis TAIR10 genome were used for Fisher’s statistical testing. Results were corrected by Yekutieli method for multiple testing, using a 0.05 threshold. GO enrichment results of significant biological pathways are presented as supplementary material and further visualized in Cytoscape (version 3.5.1) using BiNGO (Maere et al., 2005) and Enrichment Map (Merico et al., 2010) plugin.

### Basal thermotolerance assay

Arabidopsis seeds were surface-sterilized using 70% ethanol for three mins and washed with sterile water three times before sowing on ½ MS growth media plates. For the quercetin-supplemented media, quercetin (Sigma, Q4951) was dissolved in ethanol and added to the ½ MS media after autoclaving to the final concentration of 200 μM. The seeds were placed on a media with or without the addition of quercetin and incubated at 4°C in dark for 3 days to ensure synchronized germination. The plates were then transferred to the growth chamber (Percival) with 22°C under 16h / 8h of light / dark cycle (∼100 μmol m−2 s−1) with 60% air-humidity. 7 days old seedlings were subjected to heat treatment for 60 min in 45°C pre-heated growth chamber and allowed to recover for four days under standard conditions (22°C) before scoring of their survival and taking of the photographs. Plants that were still green and producing new leaves were scored as surviving.

### Measurement of water content

4-week-old soil-grown Arabidopsis plants were exposed to the same heat acclimation treatment as described above. Four days after the acclimation treatment, the acclimated and non-acclimated plants were subjected to heat stress (45°C for 90 mins). Subsequently, the aerial parts of the stressed plants were excised and weighed for fresh weight every 20 mins. Water loss rates were measured using four plants each for the acclimated and non-acclimated plants. The proportion of fresh weight loss was calculated based on the initial weight of the plant. The measurements were conducted on the laboratory bench at 21°C with 65% relative humidity.

### Stomata measurement

Arabidopsis epidermal peels were obtained from the abaxial side of the fifth rosette leaves of 4-week-old plants grown in soil which were or were not subject to the acclimation treatment as described above, and imaged with Zeiss microscope X40 objective lens. Stomatal density and aperture opening were quantified from images. Counts were made from 1mm2 regions of three to five biological replicates. Comparable regions of epidermis were scored for each rosette leaf peel in the same experiment.

### Quantitative RT-PCR Analysis

Total RNA of seedling leave tissues was extracted by using the RNeasy Mini Kit (Qiagen, USA). RNA was quantified using the NanoDrop 2000 spectrophotometer (NanoDrop Technologies, USA) and quality assessed by measuring the A260/A280 ratio. DNase digestion step was performed to remove genomic DNA from the RNA samples (Invitrogen, USA). cDNA was synthesized from 1 μg RNA of independent WT and mutant plants. The cDNA was synthesized by using Superscript I Reverse Transcriptase (RT) (Invitrogen, UK) and was used for the templates of RT-PCR and quantitative RT-PCR. Quantitative RT-PCR was performed by StepOne real-time PCR system using Fast SYBR Green Master Mix kit (Applied Biosystems, USA) to quantitate the expression level of *AGL16* and *pri-miR824* transcripts.

## Acknowledgements

The research reported in this publication was supported by funding from King Abdullah University of Science and Technology (KAUST), through both baseline support to CG and MT.

## Supplemental Figures & Legends

**Figure S1.**
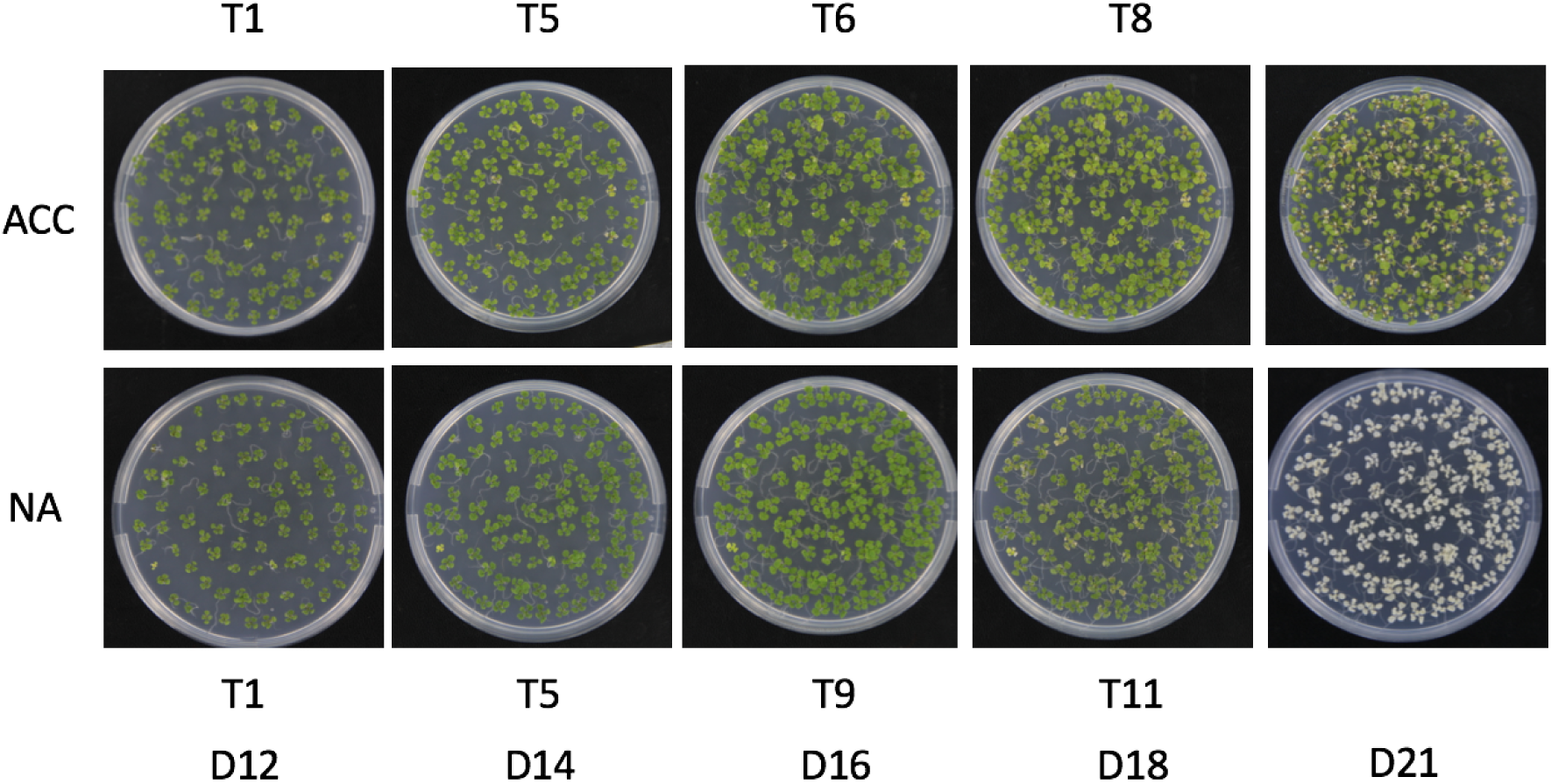
Phenotypes of acclimated and non-acclimated plants during and three days after the heat stress regime. D indicates days after sowing on the plate. T1 to T11 corresponds to the time-points in our RNA-seq experimental design (Figure 2A). ACC: acclimated. NA: non-acclimated.

**Figure S2.**
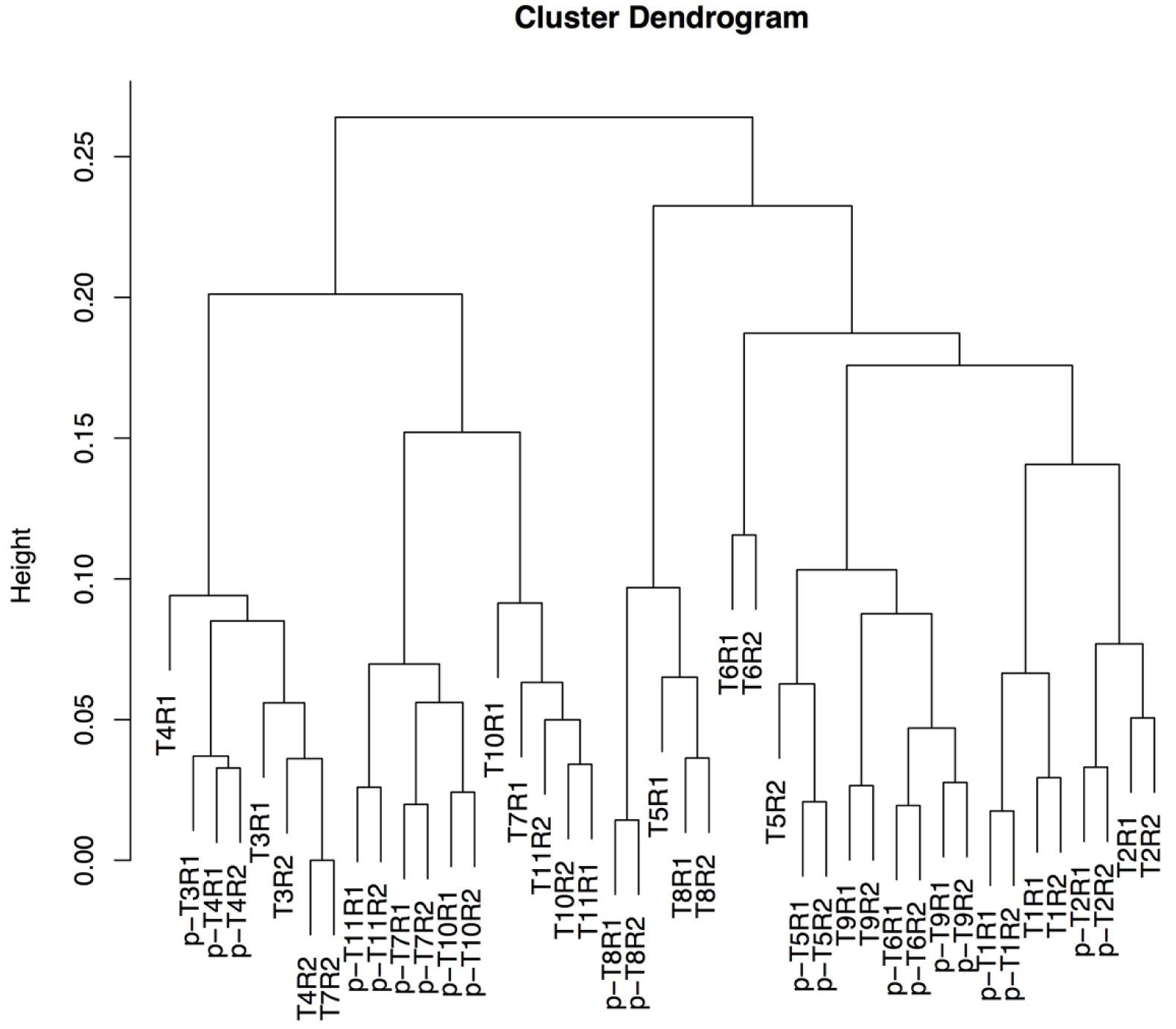
Hierarchical clustering of all the 43 samples from two independent experiments. Hierarchical clustering of samples as represented by a clustering tree. Distance between samples is measured as 1 − correlation coefficient using Ward.D agglomeration method. Samples taken at the same time points from two experiments are close to each other, suggesting that we captured the biological signals while controlling experimental errors or variations from two batches.

**Figure S3.**
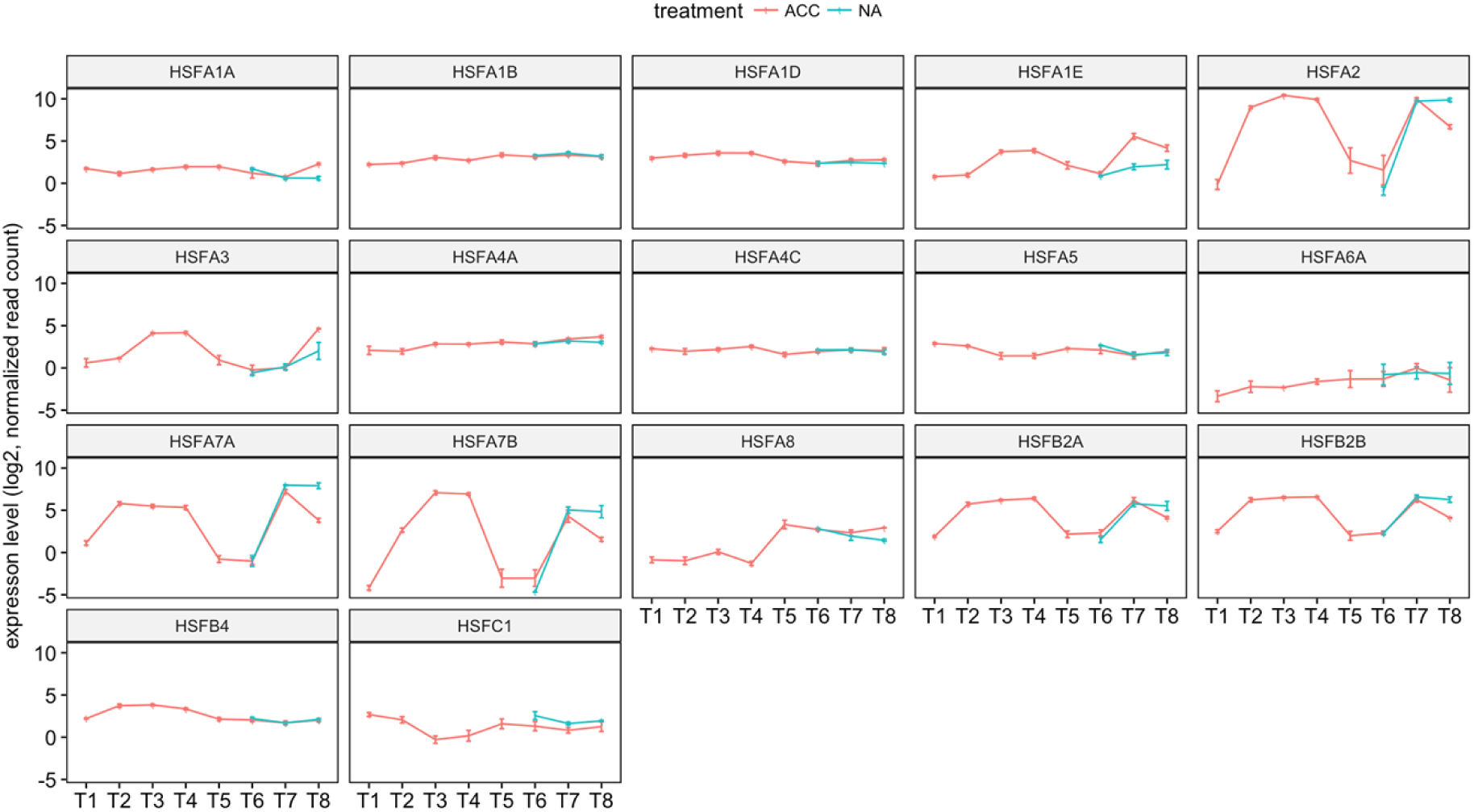
Transcripts abundance of heat shock transcription factors. Line represents the average expression of normalized read counts in log2 from 4 biological replicates, with standard error. Red line corresponds to expression pattern in acclimated plants during acclimation, before, during and after heat shock (T1 to T8). Blue line represents expression pattern in non-acclimated plants before, during and after heat shock at T9, T10 and T11 (equivalent to T6, T7 and T8).

**Figure S4.**
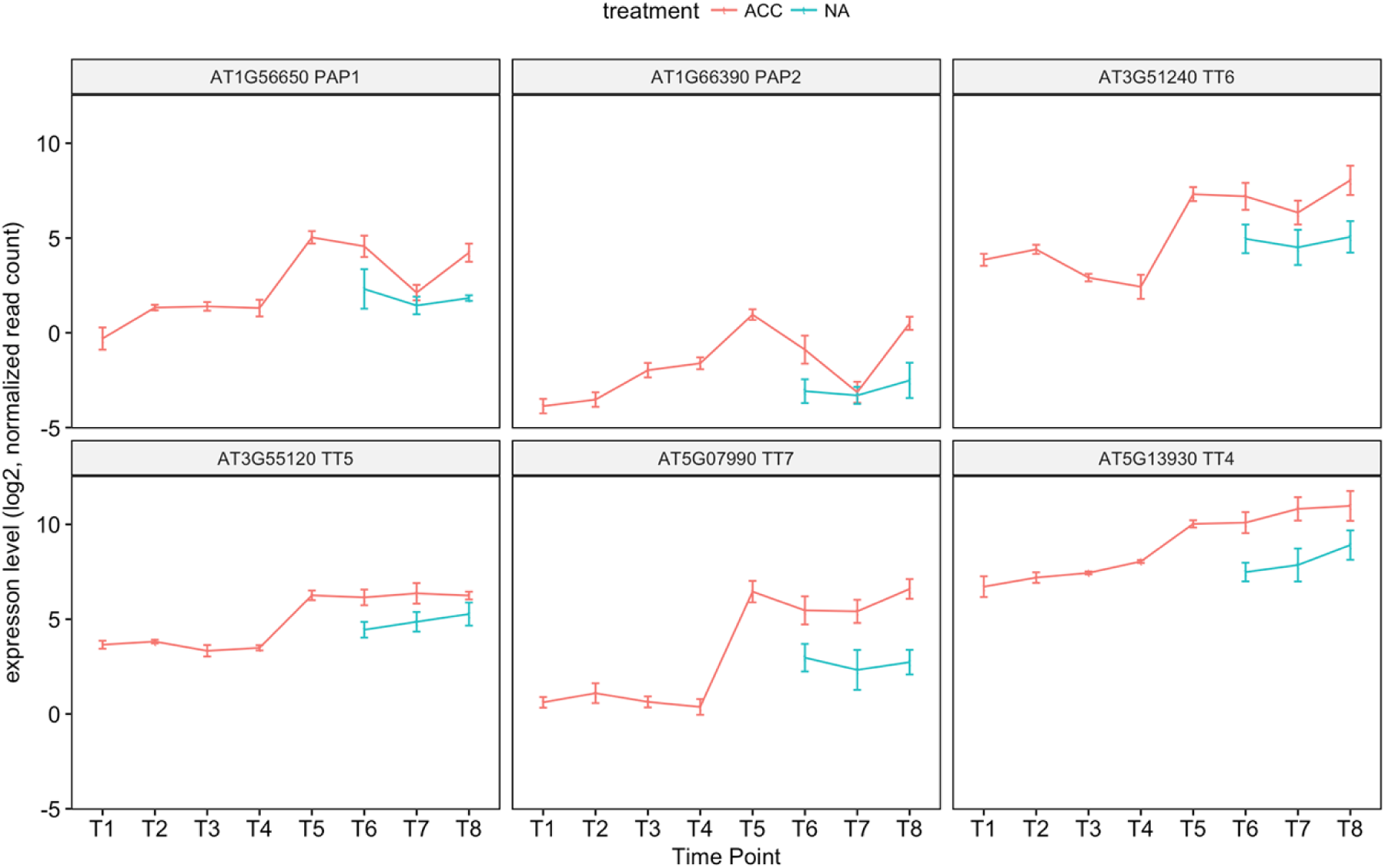
Transcripts abundance of flavonoid related genes. Each line represents the average expression of normalized read counts in log2 from 4 biological replicates, with standard error. Red line corresponds to expression pattern in acclimated plants during acclimation, before, during and after heat shock (T1 to T8). Blue line represents expression pattern in non-acclimated plants before, during and after heat shock at T9, T10 and T11 (equivalent to T6, T7 and T8).

**Figure S5.**
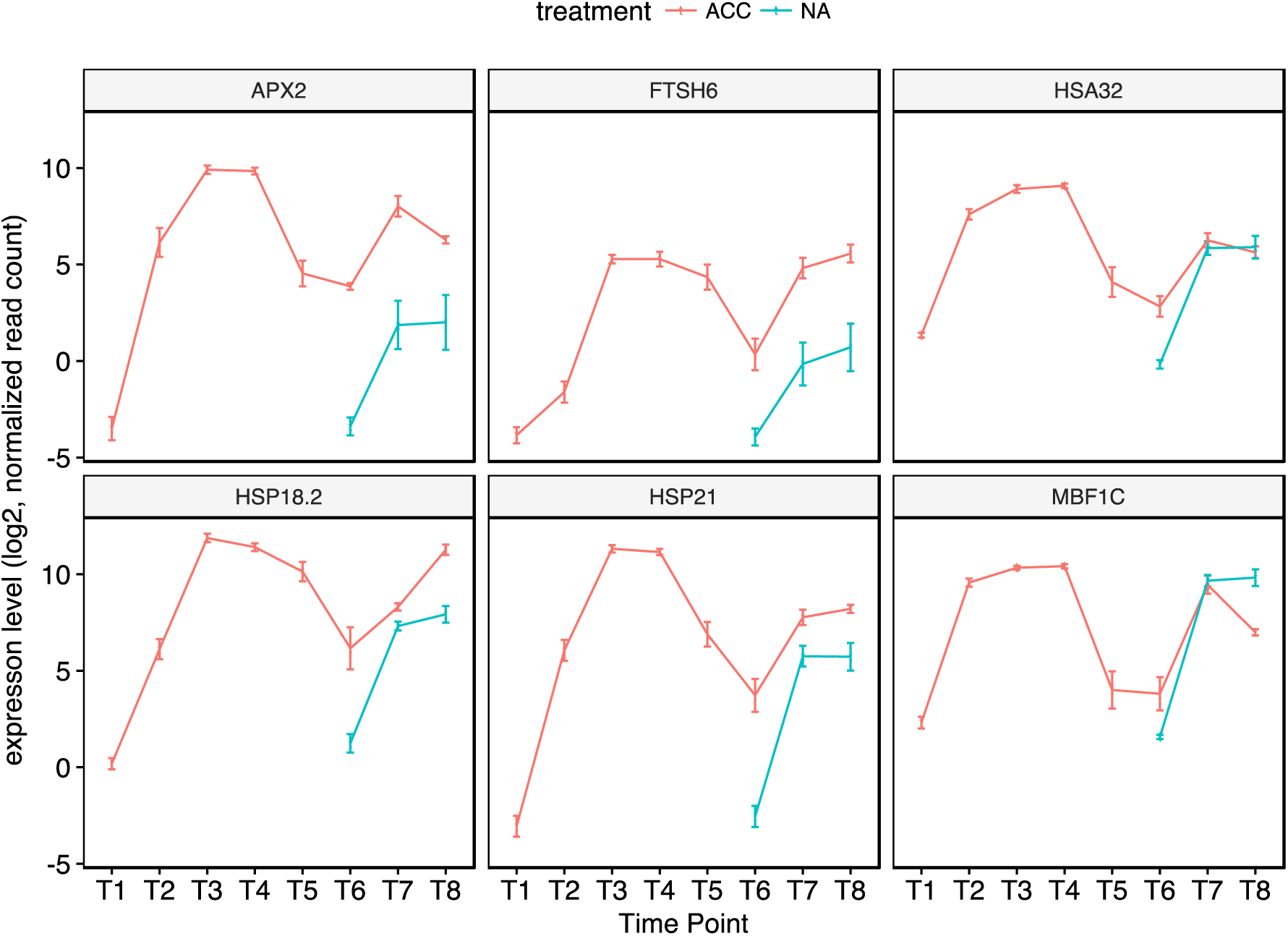
Transcripts abundance of key thermomemory genes. Each line represents the average expression of normalized read counts in log2 from 4 biological replicates, with standard error bar. Red line corresponds to expression pattern in acclimated plants during acclimation, before, during and after heat shock (T1 to T8). Blue line represents expression pattern in non-acclimated plants before, during and after heat shock at T9, T10 and T11 (equivalent to T6, T7 and T8).

## Supplemental Table legends

**Table S1.**
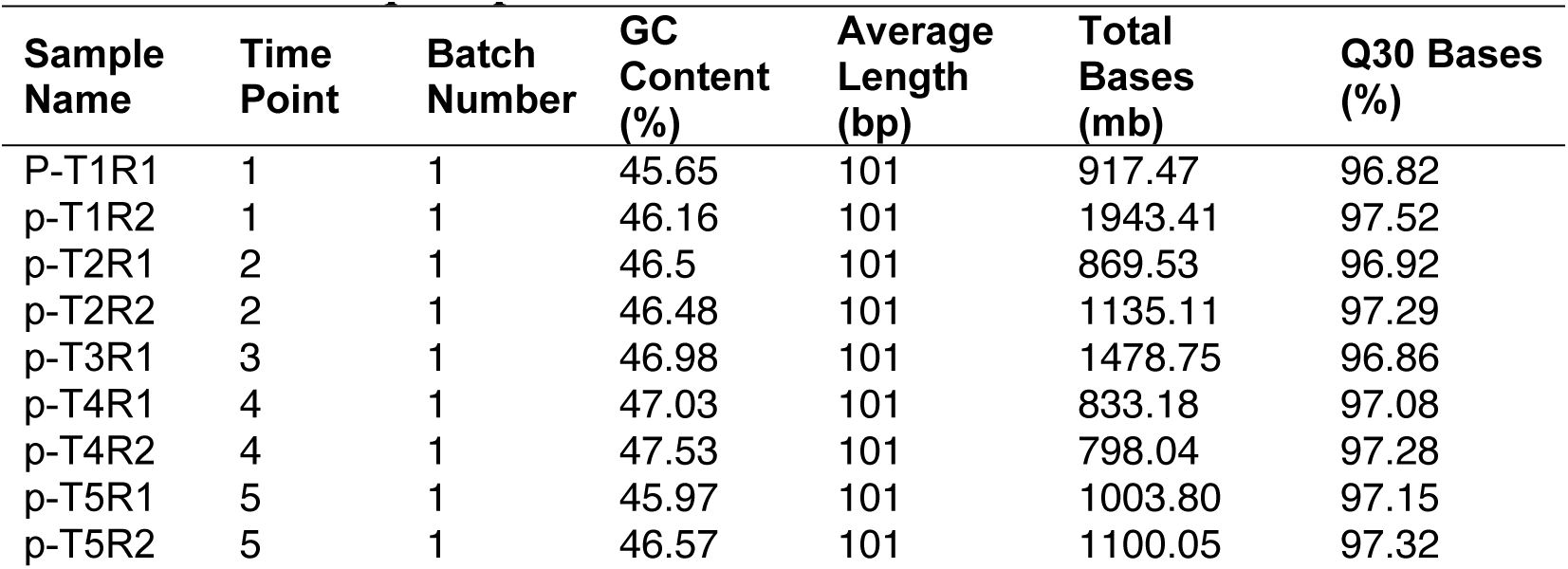
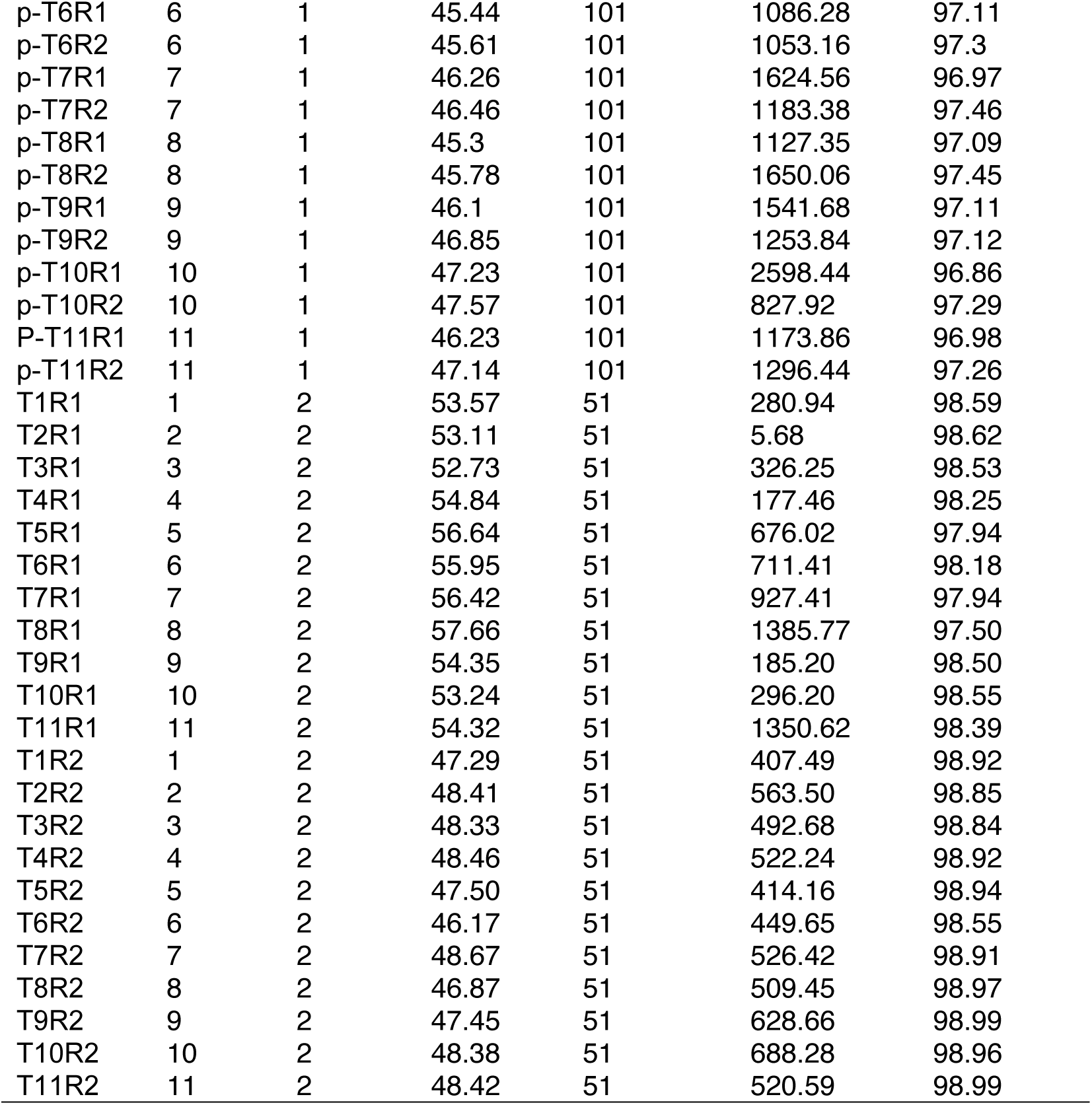
RNA-Seq sample information.

**Table S2. GO enrichment of most significant biological processes in up-regulated genes (T4 vs T1) during heat acclimation.**

**Table S3.** The R2 values for different models included in MVApp, reflecting their fit to the increase in rosette area under control and salt-stress conditions.

**Table S4.** Outliers identified using the fit of quadratic function to rosette area in MVApp. DELTA represents plant growth (mm2/day2); INTERCEPT represents the starting value of each plant. The goodness of fit of the linear model used to fit the square root transformed rosette area is represented by r_squared.

**Table S5.** List of samples identified as outliers by the 1.5*IQR method, based on all traits. A data point was considered an outlier if the sample was identified as an outlier in at least twelve traits.

**Table S6.** Correlation between individual traits measured for plants grown under salt-stress and control conditions. The correlation coefficients were calculated using curated data, with a total number of 966 samples. The outliers were identified using the 1.5*IQR method on all the measured traits; a sample qualified as an outlier if it was outlying in at least twelve traits. The data was additionally curated using the quadratic fit (samples with R2 values below 0.7 were removed).

**Table S7.** Eigen values PCA using all measured traits and curated dataset. The data were scaled prior to PCA and subset by “treatment”. PCA was performed after the outliers were removed from the curated data. The outliers were identified using the 1.5*IQR method on all the measured traits; a sample qualified as an outlier if it was outlying in at least twelve traits. The data were additionally curated using the quadratic fit (samples with R2 values below 0.7 were removed).

**Table S8.** Contributions (%) of the individual measured traits to the PCs under control and salt-stress conditions. The data were scaled prior to PCA and subset by “treatment”. PCA was performed after the outliers were removed from the curated data. The outliers were identified using the 1.5*IQR method on all the measured traits; a sample qualified as an outlier if it was outlying in at least twelve traits. The data were additionally curated using the quadratic fit (samples with R2 values below 0.7 were removed).

**Table S9.** Clustering of the phenotypes of nine Arabidopsis accessions using hierarchical clustering with the Ward method and Area, FvFm_Lss4 and NPQ_Lss4 as the major determinants. Clustering was performed after the outliers were removed from the curated and scaled data. The outliers were identified using the 1.5*IQR method on all the measured traits; a sample qualified as an outlier if it was outlying in at least twelve traits. The data were additionally curated using the quadratic fit (samples with R2 values below 0.7 were removed).

**Table S10.** Quantile regression of various traits of major interest, calculated using MVApp. The p-values from the regression analysis were calculated using MVApp for each quantile of a trait of major interest. The significant contributions are highlighted in red. The chlorophyll fluorescence traits are scored at increasing light intensities, which are marked by Lss1, Lss2, Lss3 and Lss4. The contributions of a trait of major interest to itself are not applicable (n.a.) and are marked as such in the table. The quantile regression was calculated using raw data separated by day of measurement and treatment.

**Table S11.**
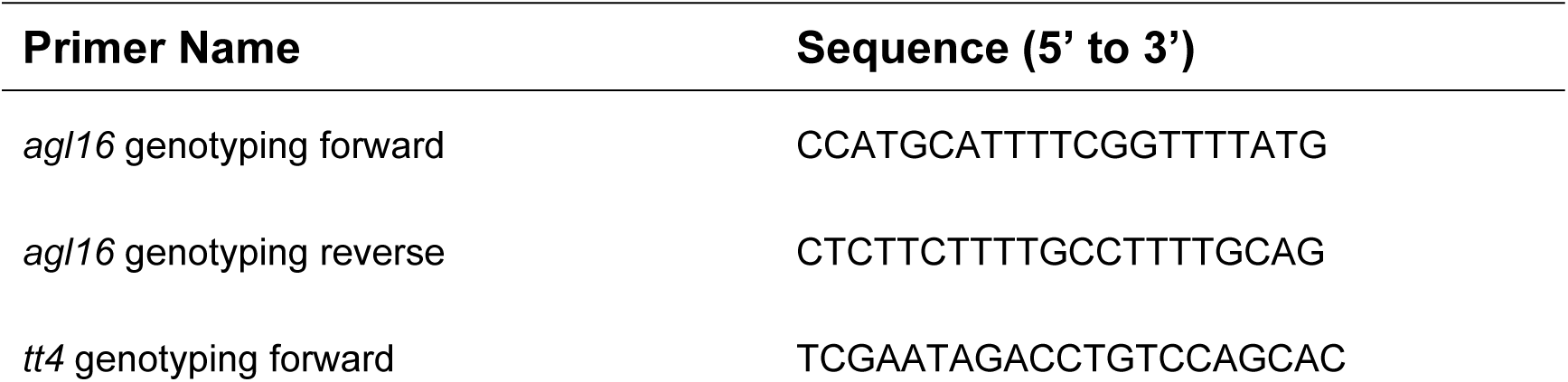
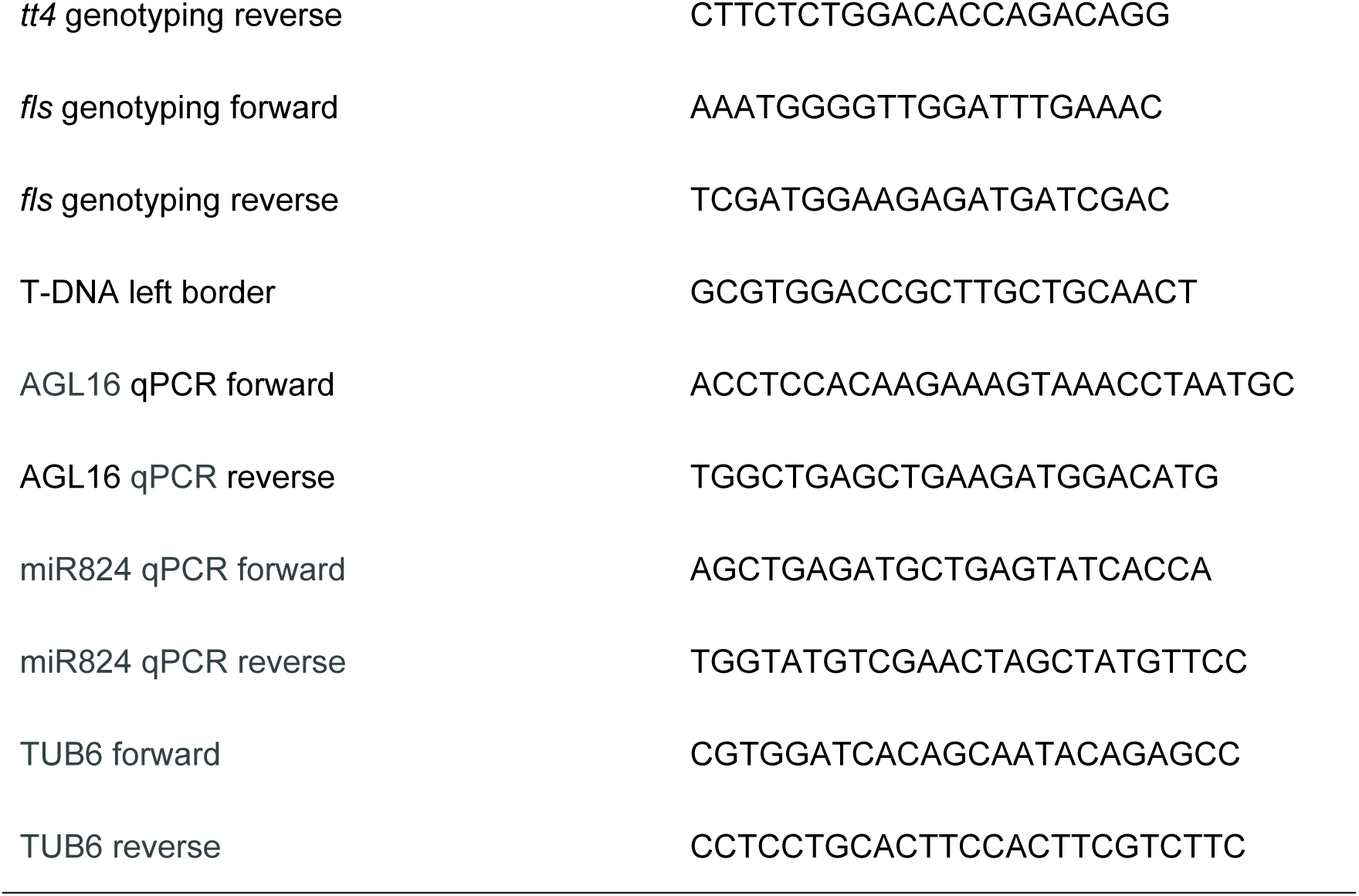
Primers used in this study.

